# Does forest heterogeneity affect mean throughfall for regenerating secondary forests on Borneo?

**DOI:** 10.1101/2022.07.14.500051

**Authors:** Nadine Keller, Ilja van Meerveld, Christopher David Philipson, Gregory P. Asner, Elia Godoong, Hamzah Tangki, Jaboury Ghazoul

## Abstract

Tropical landscape regeneration affects hydrological ecosystem functioning by regulating the amount of water that reaches the soil surface and changing soil infiltration rates. This affects the recharge and storage of water in the soil and streamflow responses. Therefore, it is important to assess how the fraction of rainfall that reaches the forest floor changes as secondary forests mature, and how forest structure affects throughfall via changes in storage capacity and evapotranspiration. Therefore, we monitored throughfall for twelve regenerating, logged-over forest plots in Sabah, Malaysian Borneo over a 7-month period and tested if inclusion of measures of forest heterogeneity improved the prediction of throughfall as a fraction of precipitation. On average across all plots, throughfall was 84% of precipitation, but was lower (as low as 74%) in plots with a longer recovery time since logging. There was a significant relationship between throughfall and tree density and basal area, as well as the Shannon Diversity Index and the coefficient of variation of the diameter at breast height, although species and structural diversity measures (Shannon Index and the coefficient of variation) did not improve model performance substantially. The overall best performing model was a linear regression with tree density. There was no relation between LiDAR-derived Top of Canopy (TCH) and mean throughfall, suggesting that this remotely sensed proxy of canopy height is not needed to estimate throughfall and more in-depth analysis of other LiDAR-products such as point clouds may be required. Our results imply that estimating throughfall in this forest type can be reliably achieved using tree density, and that this is not substantially affected by species diversity or structural heterogeneity variables, at least in the context of logged and regenerating forests in Sabah.

**Graphical abstract:** Forest of a similar size or height can have a different structure. In this study we investigate if diversity also affects the amount of throughfall for plots across a disturbance gradient.

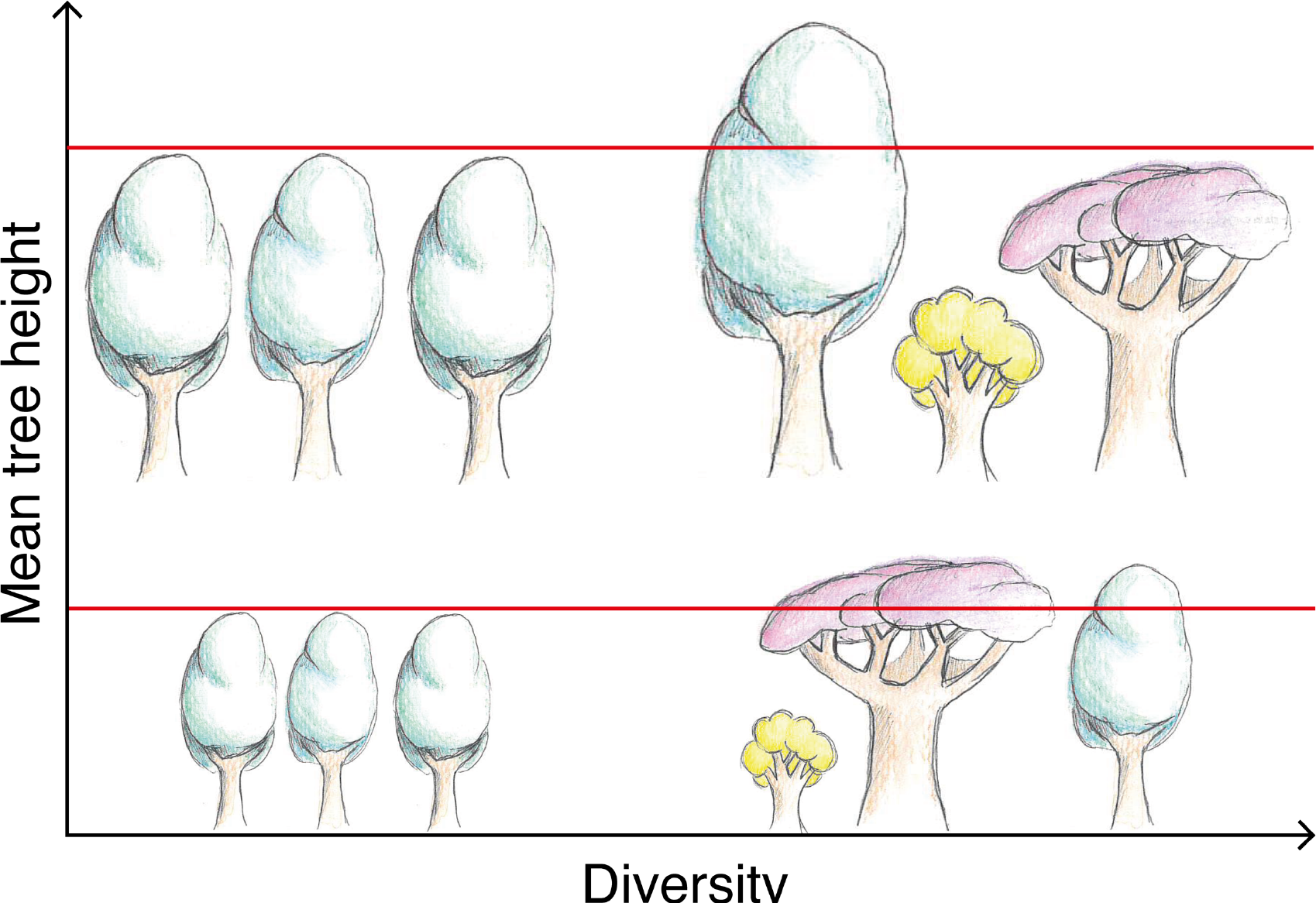

## 1. Introduction

Rates of landscape degradation remain high in the tropics. Rates of forest disturbance, mainly through deforestation and natural or anthropogenic degradation, between 1990 – 2019 were highest in Asia-Oceanian tropical moist forests, where the undisturbed area declined by 38% (i.e., by 118 Mha) (Vancutsem et al. 2021). At the same time, tropical forests are the focus of restoration efforts. Natural regeneration occurs in formerly logged forests and after abandonment of agricultural land, but can be accelerated through active forest restoration by enrichment planting and climber cutting (Chazdon 2014; Philipson et al. 2020). Most restoration projects have carbon sequestration and ecological restoration objectives (e.g., Wheeler et al. 2016), but restoring degraded forests can also provide hydrological benefits (Bruijnzeel 2004; Chazdon 2014). Soil hydraulic conductivity is higher in (actively) restored forests than on agricultural land (Ilstedt et al. 2007; Hassler et al. 2011; Lozano-Baez et al. 2019; Zwartendijk et al. 2017), which can lead to more infiltration and less overland flow and soil erosion (Bruijnzeel 2004; van Meerveld et al. 2021). Moreover, a proportion of incoming rainfall is intercepted by leaves and branches and evaporates during and after the rainfall event (i.e., interception loss). Forest stands with a higher total leaf and branch area (i.e., leaf area index (LAI)) can intercept more water, reducing the amount of water that reaches the forest floor (Ponette-González et al. 2010). Together with the changes in transpiration (Lamb 2011; Bart et al. 2021), this affects the soil water balance.

Several studies in the tropics (e.g., Dietz et al. 2006; Ponette-González et al. 2010; Zimmermann et al. 2013; Rodrigues et al. 2022) and elsewhere (Fathizadeh et al. 2017; Nadkarni and Sumera 2004; Ma et al. 2021; J. Liu et al. 2018; T. Blume et al. 2022) have investigated the relation between forest structure and throughfall or interception. These studies have mainly focused on the influence of basal area, diameter at breast height (DBH), mean tree height, LAI, tree density, or measures of biodiversity (i.e., Shannon Diversity Index; Shannon and Weaver 1949) on the amount of throughfall. For example, throughfall (the sum of amount of water that reaches the forest floor through gaps in canopy, or drips off leaves and branches) and stemflow (water that flows along tree stems to the soil) was much lower for older lowland rainforests in Panama than for young (<10 years old) secondary forests (Zimmermann et al., 2013). The ratio of the basal area of small stems (DBH: 1- 5 cm) to the total basal area was the best predictor of throughfall amount, but other measures of diversity, such as Shannon Diversity Index (Shannon and Weaver 1949), did not seem to have a significant influence on throughfall amounts (Zimmermann et al. 2013). However, there were no detectable differences among the different older forest plots (including mature secondary forests). This suggests that age effects on throughfall are only detectable for young, regenerating forests. Similarly, Ghimire et al. (2017) found a significant difference in throughfall for a young (5 – 7 years) and semi-mature lower montane secondary forest (20 years old) in Madagascar. In both studies, the younger forests were characterized by a lower leaf area index (LAI), basal area, or tree height, but also had a different forest structure and species composition.

Dietz et al. (2006) found that combining mean tree height and LAI in a multiple linear regression provided the best model to predict throughfall for four differently managed lower montane rainforest sites (natural forest, forest with small-diameter logging, forest with selective logging of large-diameter trees, and cacao agroforest) in Central Sulawesi (Indonesia). They also found a significant relation between throughfall and DBH, as well as measures related to crown size and position. Similarly, Ponette-González et al. (2010) documented a significant relation between throughfall and LAI and basal area for a Mexican montane landscape consisting of montane cloud forest, coffee agroforestry, and cleared areas and showed that the best regression model combined information on the basal area and mean minimum tree height. However, LAI and basal area were strongly correlated with each other. Studies conducted outside tropical regions have highlighted the importance of complex canopies (J. Liu et al. 2018), or the presence of epiphytes (Nadkarni and Sumera 2004) on the throughfall amount. The varying importance of the different predictors for throughfall amount may reflect regional differences in climate or forest composition (Ghimire et al. 2017) and highlights the importance of obtaining local models to estimate throughfall based on forest characteristics.

Borneo is home to one of the most valuable tropical forests in terms of biodiversity (Myers et al. 2000) and sequestered carbon (Ioki et al. 2014), but is also experiencing very high deforestation rates (Laurance and Edwards 2014): the forest area declined by 33% between 1973 – 2015 (Gaveau et al. 2016). Several previous studies have investigated throughfall in Borneo, but they all focused on quantifying throughfall amounts and did not explore the influence of forest characteristics on mean throughfall (Sinun et al. 1992; Dykes 1997; Asdak et al. 1998; Chappell et al. 2001; Vernimmen et al. 2007). For example, Chappell et al. (2001) found that mean throughfall for selectively logged, moderately impacted lowland dipterocarp forests (84.1 ± 6.9% of precipitation) was high compared to moderately to very disturbed forests (79.6 ± 6.9% of precipitation). Asdak et al. (1998), on the other hand, measured throughfall values of 89% for pristine forests and 94% for logged forests.

The relation between throughfall and forest characteristic at the stand scale can be used to estimate throughfall or interception losses for larger areas or regions. However, this requires data on the structure of the forests, which is time-consuming to collect. Remote sensing data derived from satellites (e.g., Nieschulze et al. 2009; Hassan et al. 2017) and airborne LiDAR data (Teske and Thistle 2004; Roth et al. 2007) can provide useful information on forest characteristics. Nieschulze et al. (2009), for instance, successfully predicted throughfall with geostatistical and spectral parameters for four different land-use types (natural forest, forests with selective logging of small or large timbers, and cacao agroforest) in central Sulawesi (Indonesia) using high-resolution optical satellite imaginary. Roth et al. (2007) emphasized the opportunity to map canopy architecture in 3D using LiDAR, and the potential to improve rainfall interception estimates over a large spatial scale.

However, most previous studies on throughfall in regenerating tropical rainforests forests used only mean values of forest structure and ignored measures of forest heterogeneity. This is problematic as a uniform, plantation-like forest can have the same mean tree height as a diverse, highly complex primary forest (see graphical abstract). A linear model based on tree height would then predict the same amount of throughfall in both forests, even though interception is expected to be higher (and throughfall lower) for a heterogeneous forest with a range of leaf, tree, and crown shapes and sizes because structural complexity implies a more complete use of space in the forest (e.g., a more layered canopy structure). The more complete use of the space increases the likelihood of raindrops being intercepted, while leaf heterogeneity could increase the retention of water on leaf surfaces in different ways, thereby further slowing vertical water flow through the canopy and increasing the possibility of evaporation during the event. Consequently, inclusion of measures of diversity to reflect the heterogeneity of forest and tree structure may improve throughfall predictions at the stand scale. Indeed, Nadkarni and Sumera (2004) report a strong relationship between throughfall and the diversity of structural elements for an old-growth coniferous forest in Washington State (USA). Similarly, J. Liu et al. (2018) found lower throughfall in forests with a higher vertical heterogeneity or higher structural complexity in Beijing (China).

We, therefore, measured throughfall in 12 regenerating forest plots in Borneo and compared the performance of statistical models using only the mean values of structural forest characteristics with models that additionally included information on forest heterogeneity, such as the variance around the means, and tree diversity. We compared the relationships between mean throughfall and (structural) forest characteristics using both ground-measured data and data derived from airborne LiDAR. More specifically, we aimed to answer the following questions:

1. What is the relation between throughfall and structural forest characteristics for regenerating secondary forests in Sabah, Malaysian Borneo?
2. Does the inclusion of forest heterogeneity measures improve the prediction of throughfall compared to models based on forest structural characteristics (such as tree size or density) alone?
3. Can the mean throughfall be predicted with LiDAR-derived top of canopy height (TCH), and can model performance be improved by including information on the variation in TCH?

## 2. Study site description

### 2.1. General location and climate

This study was conducted in Southeast Sabah, Malaysian Borneo, which climate is typical for equatorial rainforests. The average annual precipitation is approximately 2700 mm/y, and is distributed relatively uniformly throughout the year (Walsh and Newbery 1999). The monthly mean temperature is 27°C (± 1.9°C) (Walsh and Newbery 1999). Relative humidity is generally very high. It is close to saturation (100%) at 8.00 am and lowest in the afternoon (∼72% at 2pm) (Walsh and Newbery 1999). High intensity rainfall events occur relatively frequently (intensity > 50 mm/hr (maximum 5 min intensity) with return-period of 23 days, and > 100 mm/hr (maximum 5 min intensity) with return-period of 140 days) (Sherlock 1997). While the climate is influenced by El Niño-Southern Oscillation events that occur irregularly every few years and cause unusually dry periods lasting several months (Walsh and Newbery 1999), no such event occurred during our 7-month measurement period.

### 2.2. Description of the forest in the two study areas

The study was conducted in two different areas within the Ulu Segama-Malua and Kinabatangan Districts: Kuamut in Kinabatangan, and INFAPRO in Ulu Segama-Malua. Working in the two different areas allowed us to measure throughfall for stands with different characteristics. The two areas are located 25 km apart, differ by less than 170 m in elevation, are ecologically similar and border the Danum Valley Conservation Area. Soil type is similar for both areas as well and classified as orthic acrisol (Hector et al. 2011; Saner et al. 2012; H.- P. Blume et al. 2010).

The forests in both areas are classified as lowland dipterocarp dominated. The areas differ in their logging and restoration history. The logging history of Kuamut area is not well documented, so we rely on information from the neighbouring Malua forest reserve (coordinates of entrance gate: N 5° 25’ 33; E 117° 1’ 58). The Malua forest reserve was selectively logged twice, once in the 1980s, and again between 2003 and 2007 (Reynolds et al. 2011). The forest has re-grown naturally (other than the Sabah Biodiversity Experiment, there has not been any active restoration) (Saner et al. 2012; Reynolds et al. 2011; Philipson et al. 2014) but the logging activities have led to a generally degraded forest with different levels of disturbance (Jucker et al. 2018). The INFAPRO area (N 4° 58’ 52; E 117° 51’ 25) was selectively logged between 1972 and 1993, and was subject to active restoration (enrichment planting of seedlings and climber-cutting) in some areas between 1993 and 2004. However, other areas were left for natural regeneration (Philipson et al. 2020).

### 2.3. Plot selection

We selected twelve study plots: nine in Kuamut and three in the INFAPRO area. Each 0.28 ha circular forest plot had a 60 m diameter. The plot locations were chosen to cover a range of forest characteristics, in particular canopy height and species diversity. Potential locations of interest were identified using the Top of Canopy Height (TCH) maps (derived by airborne light detection and ranging (LIDAR); Asner et al. (2017)). After selection of the plot for inclusion in the study, the number of species, tree height, etc. were measured (see section 3.1.1). We selected plots with similar vegetation structure that did not have a sudden transition to more open or closed forest for at least 30 meters outside the plot.

## 1. 3. Methods

### 3.1. Forest characteristics

#### 3.1.1. Ground-derived field measurements

In each plot, we determined for all tree or liana stems with DBH larger than 10 cm, the diameter at breast height (DBH), height, and species. DBH was measured with a measurement tape; tree height with an ultrasound rangefinder (Vertex IV; Haglöf). If buttresses were present, DBH measurements were taken ten cm above them. A tapering model (with a fixed parameter value, as suggested by Cushman et al. (2014)) was used together with these two measurements to estimate the DBH at the standard height of 1.3 m (Metcalf et al. 2009; Jucker et al. 2018). Trees were identified to family, genera, and species (if possible) by examining their leaves. Lianas were treated as one group (“liana”). As only 1.6% (20 individuals out of 1276) of all recorded stems belong to the liana group, we simply us the term “trees” in this study. Moreover, trees for which the canopy was not visible or were defoliated, and samples that could not be matched to a species, were handled as unidentified, and treated as unidentified (i.e., NAs). In summary, 60% of all trees were identified at the species level, 31% at the genus level, and 7% remained unidentified (Table 1).

**Table 1:** Characteristics of the 12 forest plots (9 in Kuamut, indicated by a plot-name starting with "*K*", and 3 in INFAPRO, indicated by plot-name starting with "*I*"). *No. stems* is the number of stems with a diameter at breast height (DBH)>10 cm (i.e., number of trees and lianas combined) per plot, whereby *No. lianas* is the number of lianas per plot. *Percentage identified trees on species / genus - level* represents the percentage of trees with a DBH>10 cm that could be identified at the species- or genus-level. *Tree density, mean tree height*, *coefficient of variation (CV*) *of DBH, and Shannon Diversity Index* are based on field measurements. *Mean top of canopy height* and *CV of top of canopy height* are based on LIDAR measurements. Please note that LiDAR-data was not available for the INFAPRO plots (as indicated by the NA). *Mean throughfall* is the arithmetic mean of the average throughfall in percentage of rainfall for each plot. *No. throughfall measurements* give the number of measurement dates that were included in the calculation of the mean throughfall.

| Plot name | coordinate | No. stems (-) | No. lianas (-) | Percentage identified trees on species / genus - level (%) | Tree density (#trees / ha) | Mean tree height (m) | Shannon Diversity Index (-) | CV of DBH (-) | Mean top of canopy height (m) | CV in top of canopy height (-) | Mean throughfall (%) | No. throughfall measurements (-) |
| --- | --- | --- | --- | --- | --- | --- | --- | --- | --- | --- | --- | --- |
| K1 | 05°03'24.0''N<br>117°35'09.3''E | 115 | 2 | 56 / 36 | 406.7 | 13.4 | 3.50 | 1.07 | 29.9 | 0.51 | 82 | 31 |
| K2 | 05°03'26.1''N<br>117°35'12.6''E | 106 | 2 | 68 / 23 | 374.9 | 15.0 | 2.96 | 0.67 | 15.4 | 0.49 | 88 | 31 |
| K3 | 05°03'21.3''N<br>117°35'11.1''E | 62 | 3 | 69 / 31 | 219.3 | 19.2 | 3.29 | 0.93 | 26.7 | 0.51 | 89 | 31 |
| K4 | 05°03'20.4''N<br>117°35'12.0''E | 109 | 4 | 46 / 52 | 385.5 | 19.5 | 3.67 | 0.82 | 29.1 | 0.42 | 88 | 31 |
| K5 | 05°03'22.4''N<br>117°35'12.8''E | 77 | 1 | 61 / 32 | 272.3 | 17.6 | 3.48 | 0.80 | 32.6 | 0.54 | 86 | 31 |
| K6 | 05°03'27.8''N<br>117°35'09.9''E | 128 | 1 | 78 / 21 | 452.7 | 13.2 | 3.40 | 0.72 | 22.3 | 0.43 | 82 | 31 |
| K7 | 05°03'42.5''N<br>117°35'41.3''E | 75 | 0 | 96 / 3 | 265.3 | 17.6 | 0.86 | 0.56 | 15.3 | 0.59 | 89 | 29 |
| K8 | 5°03'22.6''N<br>117°34'59.0''E | 125 | 5 | 33 / 43 | 442.1 | 17.3 | 3.17 | 0.91 | 31.4 | 0.48 | 81 | 32 |
| K9 | 5°03'41.0''N<br>117°35'11.1''E | 87 | 3 | 77 / 8 | 307.7 | 16.1 | 3.14 | 0.53 | 15.0 | 0.51 | 89 | 30 |
| I1 | 04°57'80.8''N<br>117°50'11.7''E | 131 | 0 | 67 / 24 | 463.3 | 18.3 | 3.80 | 0.92 | - | - | 74 | 28 |
| I2 | 04°59'58.6''N<br>117°49'27.2''E | 135 | 0 | 60 / 37 | 477.5 | 20.2 | 3.81 | 1.00 | - | - | 78 | 28 |
| I3 | 04°59'89.1''N<br>117°49'36.4''E | 129 | 0 | 43 / 51 | 445.6 | 18.6 | 3.53 | 0.81 | - | - | 84 | 28 |

Based on these plot measurements, the following forest characteristics were calculated: tree density, mean height of all trees, mean height for the 5% tallest trees, mean DBH, and total basal area. In addition, we calculated the Shannon Diversity Index and the Simpson Diversity Index (Shannon and Weaver 1949; Simpson 1949) using the R package *vegan* (Oksanen et al. 2020) in R studio (version 1.4.1106) (R Core Team 2021). Finally, we determined the coefficient of variation (i.e., standard deviation divided by the mean value) of tree height and DBH as additional diversity indicator (see Table S1 for more detailed documentation of the calculations).

#### 3.1.2. LiDAR-derived Top of Canopy (TCH)

In addition to the field data, we also used Top of Canopy Height (TCH) maps based on airborne light detection and ranging (LiDAR) data from Asner et al. (2017). The TCH maps were only available for the plots located in the Kuamut region (n = 9), and not for the plots at INFAPRO region (n = 3). The LiDAR data were collected by the Global Airborne Observatory (GAO; formerly known as the Carnegie Airborne Observatory; Asner et al. (2012)) in 2016 (i.e., two years before our study). The TCH maps were derived from analysis of the laser point cloud data following removal of ground returns, resulting in a TCH map with 1 m spatial resolution, where the top of canopy is shown independent of the elevation where trees are growing.

The TCH represents a similar forest characteristic as the (ground-derived) mean tree height, but differs in the way that they are calculated. For the calculation of mean tree height, each tree within the forest plot has the same weight, regardless of the tree size or crown diameter. The mean TCH only considers the height of the uppermost canopy layer, and represents the contribution of individual trees in the upper most canopy to the mean TCH based on the size of the tree crown (as trees with a larger crown cover a larger area of the forest plot).

The GPS used to determine the location of the plots (Garmin GPSMAP 64s) had an accuracy of only ±7 m when used in the forest. We accounted for this low accuracy by creating a buffer on the TCH map with a 7 m radius around the center coordinates and then randomly chose 1000 alternative plot middle points from inside this buffer. For each of these 1000 points per forest plot we calculated for the area within the 30 m radius from the center point, the mean TCH, and the coefficient of variation of TCH (CV TCH) (see Table S1). We used the average of these 1000 values to obtain one value for the mean and the CV of TCH per plot.

### 3.2. Throughfall measurements

In each of the twelve plots, we installed 50 rainfall gauges on a regular (7 by 7) square grid, with 6 m between the gauges in all direction (see Supplementary material 1). The 50^th^ gauge was installed 6 m to the north of the middle grid line (see Figure S2). All gauges had a 9.0 cm diameter circular opening and were attached to a wooden poles at 1 m above the ground to avoid ground-splash (cf. Ghimire et al. 2017; see Figure S3).

The throughfall gauges were emptied each morning after a rainfall event between March and October 2018. The daily sampling meant that if multiple events occurred on one day, they were considered as one event (see Supplementary material S2). If data collection was not possible in the morning after a rainfall event (e.g., due to inaccessibility of field sites due to flooding, or the presence of elephants), we measured the amount of water in the gauges the next morning, or the day after. If the gauges could not be emptied within 72 hours after a rainfall event, they were emptied without recording the amount of rainfall/throughfall. In case rainfall occurred during data collection, the collection was aborted, and none of the collected data were used. In total, data were collected for 30 to 34 events at each plot. However, some of these data points had later to be dismissed (see section 3.3, and Table 1 for final number of data points per forest plot). We assume that the average amount of water measured in the 50 gauges per plot represents the average throughfall for that measurement date.

The throughfall gauges were moved by 1 meter three times during the study period: 1 m to the north after 40 days, 1 m to the east after 80 days, and 2 m to the west (i.e., 1 m to the west from second position) after 120 days (see Figure S2). This was done to incorporate more of the spatial variation in throughfall in the measurements (i.e., inclusion or exclusion of so-called drip- points) and, thus, to obtain a more accurate estimate of the average throughfall.

### 3.3. Rainfall measurements

Five reference rainfall stations were established in the Kuamut region, and two in the INFAPRO area. These rain gauges were installed in open areas with short vegetation (i.e., grass, ferns) and at least 10 m from the forest edge. At each rainfall station, we installed four gauges (similar to the ones used for the throughfall measurements) in a 1 x 1 m area. We used the average rainfall from the four gauges to determine the incoming rainfall for each rainfall station.

For half of the experimental plots, a reference rainfall station was located within 50 m from the plot and the average of the measured rainfall could be used for the incoming rainfall. For the other plots, we used inverse distance weighting with reference rainfall from the two or three nearest stations to determine the incoming rainfall. The longest distance between a plot and the nearest reference rainfall station was 536 m (average: 212 m). Although plots K3 and K9 were located within 50 m of a rainfall station, we had to use inverse distance weighting for a few measurement dates (n=2 for K3, and n=11 for K9) because the data for the nearest reference rainfall station was not available due to a delayed set-up or destruction by elephants.

### 3.4. Data analysis

#### 3.4.1. Mean Throughfall

We did not measure stemflow because studies in Northern Borneo (Sabah and Sarawak) indicated that stemflow was only 1.9% and 3.5% of gross precipitation, respectively (Manfroi et al. 2004; Sinun et al. 1992). We, therefore, regarded the effect of stemflow to be negligible and consider the difference between the reference rainfall and average throughfall to be the interception loss.

We divided the average throughfall by the incoming (or reference) rainfall to obtain a throughfall percentage for each measurement date. Average throughfall values were occasionally greater than the incoming rainfall (i.e., throughfall > 100%). The ten (=2.7% of all daily measurements at all the plots) measurements yielding an average throughfall greater than 120% were assumed to be due to errors in the measurements, particularly errors in the incoming rainfall, and were therefore excluded from the analyses. Spatial variation in rainfall as well as interaction with wildlife that caused the gauges to not be perfectly leveled may have caused an underestimation of the incoming rainfall as well (see Supplementary material 3 for additional information). Average throughfall values between 100 and 120% were assumed to be realistic and due to 1) small-scale differences in rainfall, 2) condensation on leaves and drip into the throughfall gauges, or 3) evaporation from reference rainfall gauges (see also discussion section 5.1).

We checked for the occurrence of a breakpoint in the relation between the average throughfall (as a percent of reference rainfall) and the incoming rainfall using the R-package *segmented* (Muggeo 2008) to divide the events into small events that did not saturate the canopy and large events that fill the canopy storage capacity (cf., the canopy storage capacity *S* in the Gash model (Gash and Morton 1978); see section 4.2). However, this did not yield reasonable results as the breakpoint value given by the *segmented* R-package (Muggeo 2008) was unreasonable large (e.g., at a reference rainfall size of 90 mm) for half of the forest plots. Moreover, splitting the data points at an arbitrary smaller reference precipitation (e.g., 15 mm cf. Brasil et al. (2018) or Molina-Sanchis et al. (2016), or 40 mm cf. Brasil et al. (2022)) or a reasonable value of the canopy storage capacity (e.g., the 1.2 mm reported by Ghimire et al. (2017) for a semi-mature secondary forest) would have led to a highly imbalanced number of data points for each size category. For most of the 12 forest plots, only very few data points (n ≤ 5) would fall in the small rainfall size category. Therefore, we decided to use the overall arithmetic mean of the average throughfall (as percent of reference rainfall) for all events in the analysis with the plot characteristics. Thus, the mean throughfall for a forest plot is the average of all the plot average throughfall values (based on the 50 individual measurements) for all the measurement dates (i.e., arithmetic mean). The arithmetic mean throughfall is highly correlated with the mean throughfall weighted by incoming rainfall (R^2^: 0.90), but the arithmetic mean throughfall value is smaller than the weighted mean throughfall (mean difference: 4%, range: 1-8%; Figure S6).

#### 3.4.2. Relation between forest characteristics and mean throughfall

We used the *corrplot* package (Wei and Simko, 2017) to create a correlation matrix to identify the correlation between the different structural forest characteristics and diversity indicators. We investigated the relation between the mean throughfall (as percentage of rainfall) and each of the structural forest characteristics (either field- or LIDAR-derived) and diversity indicators using both Pearson (linear) and Spearman rank correlation. In addition, we used linear models to examine the relationship between mean throughfall and a structural forest characteristic in combination with a diversity indicator. Based on the correlations between the structural forest characteristics and the diversity indicators (see section 4.1), we selected two key characteristics as explanatory variables for the mean throughfall in the linear models: tree density and mean tree height. As a measure of variability or biodiversity, we chose the Shannon Diversity Index because it takes species richness, as well as the relative abundance of species into account. As an alternative diversity indicator, we chose the coefficient of variation (CV) in DBH, because of it its similarity to the best predictor (i.e., ratio of basal area of small stems (DBH < 5 cm and > 1 cm) to the total basal area) of the variation in throughfall amount found by Zimmermann et al. (2013). For the LiDAR-derived structural forest traits and diversity measures, we chose the mean top of canopy height (TCH), and the coefficient of variation (CV) in TCH height as the main measures of diversity because of their similarity to the ground-derived measurements of tree height.

For the multiple linear regression, we first tested if there was a significant interaction effect between the explanatory variables (i.e., the forests structural characteristic and diversity measure). If there was no significant interaction effect, we pursued model simplification by excluding the interaction terms. As we aimed to describe the increase/decrease in model performance through the inclusion of diversity measures, we added each diversity measure individually, instead of using them all in the models at once. The performances of the models with and without diversity measures were compared using the Akaike Information Criterion (AIC). The model with the lowest AIC-value was considered the best model, if the difference in AIC-units is larger than two, otherwise the simplest model was chosen.

We ensured that model assumptions were met by looking at diagnostic plots, and inspecting Cook’s distance. The Cook’s distance assessment suggested that one plot (K7) has potentially a large influence (i.e., high leverage) on the linear models including the Shannon Diversity Index. We therefore report model results with and without including data from this particular plot. Similarly, correlation tests including only the Shannon Diversity Index were conducted for the full dataset (n=12) and without this plot (n=11).

Using AIC for comparison of model performance is only possible when the models are calculated with the same datapoints. Consequently, to allow comparability of ground-derived and LiDAR-derived data (which was only available for a subset of forest plots), the models with ground-derived data were also calculated using only the data of plots in Kuamut. Lastly, we calculated the null-model (mean throughfall ∼ 1) in order to verify that the described models actually explain more variation than is simply explained by chance.

## 4. Results

### 4.1. Correlation of different forest characteristics

Tree density varied between 219 and 478 individuals per ha, mean tree height between 13 m and 20 m, and the Shannon Diversity Index between 0.86 and 3.81 (Table 1). As expected, many of the forest characteristics and diversity measures were highly correlated (Figure 1, Table S2). For example, tree density was correlated with basal area; the average tree height was correlated with the height of the tallest 5% of the trees and DBH. However, neither the Shannon Diversity Index, nor the coefficient of variation (CV) of DBH were correlated to tree density or tree height (Figure 1). Similarly, the mean top of canopy height (TCH) was not correlated to the coefficient of variation (CV) of TCH (Figure 1, Table S2). As a result, the selected forest plots cover a spectrum of tree sizes and diversity or variability (Figure 2).

**Figure 1:**
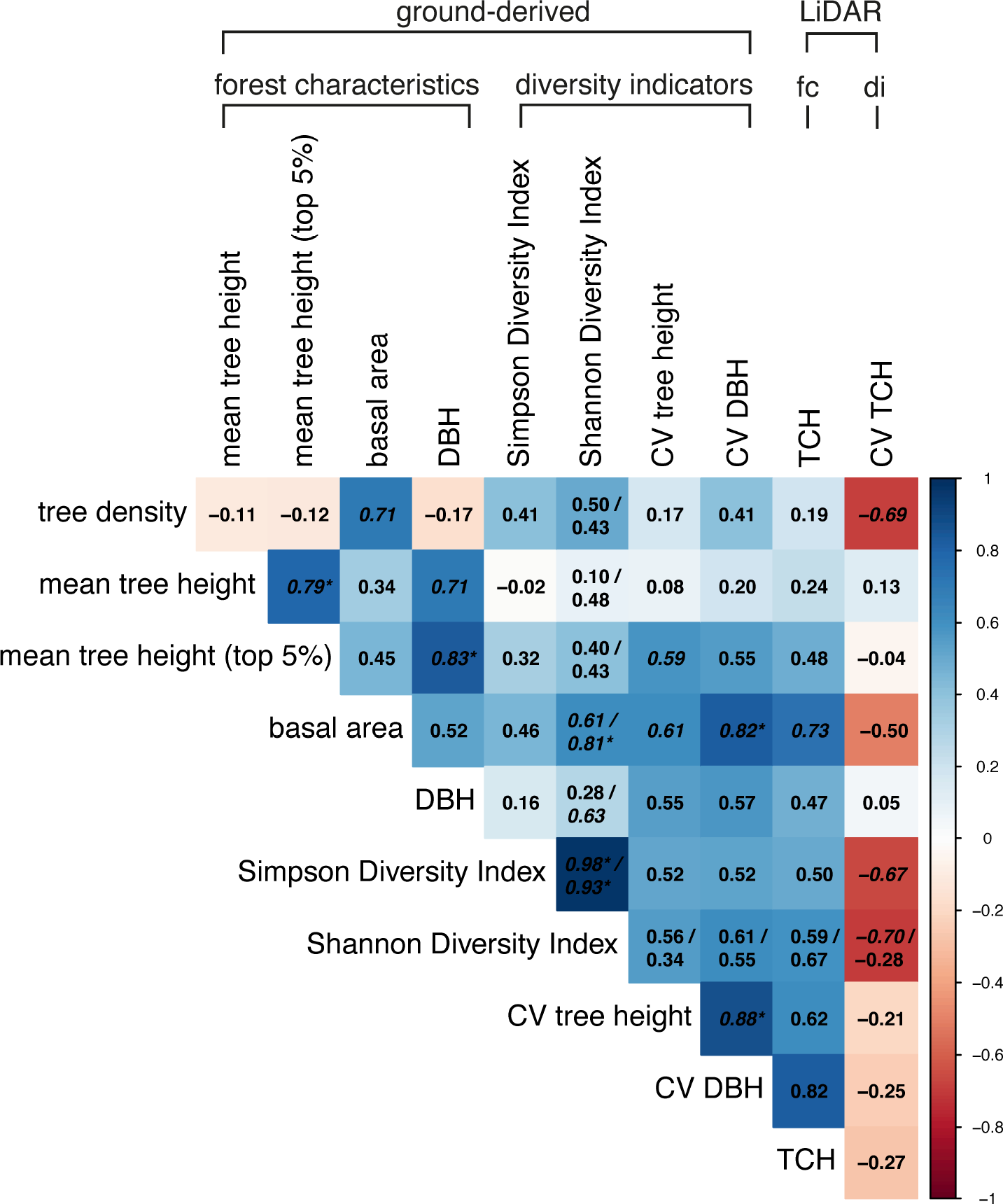
Pearson correlation coefficients between all ground-derived structural forest characteristics (fc) (tree density, mean tree height, mean tree height top 5% tallest trees, basal area, DBH) and diversity indicators (di) (Simpson Diversity Index, Shannon Diversity Index, CV of tree height), as well as the LiDAR-derived structural forest characteristic (top of canopy height; TCH) and the CV of TCH). Significant (p<0.05) correlations are italicized; correlations with p<0.01 are additionally marked with a *. Correlation values for Shannon Diversity Index are reported for two subsets of data (n=12 / n = 11). Note that the color of the tiles refers to the correlation tests with the full subset (n=12). See Figure S1 and Table S2 for the Spearman rank correlation results.

**Figure 2:**
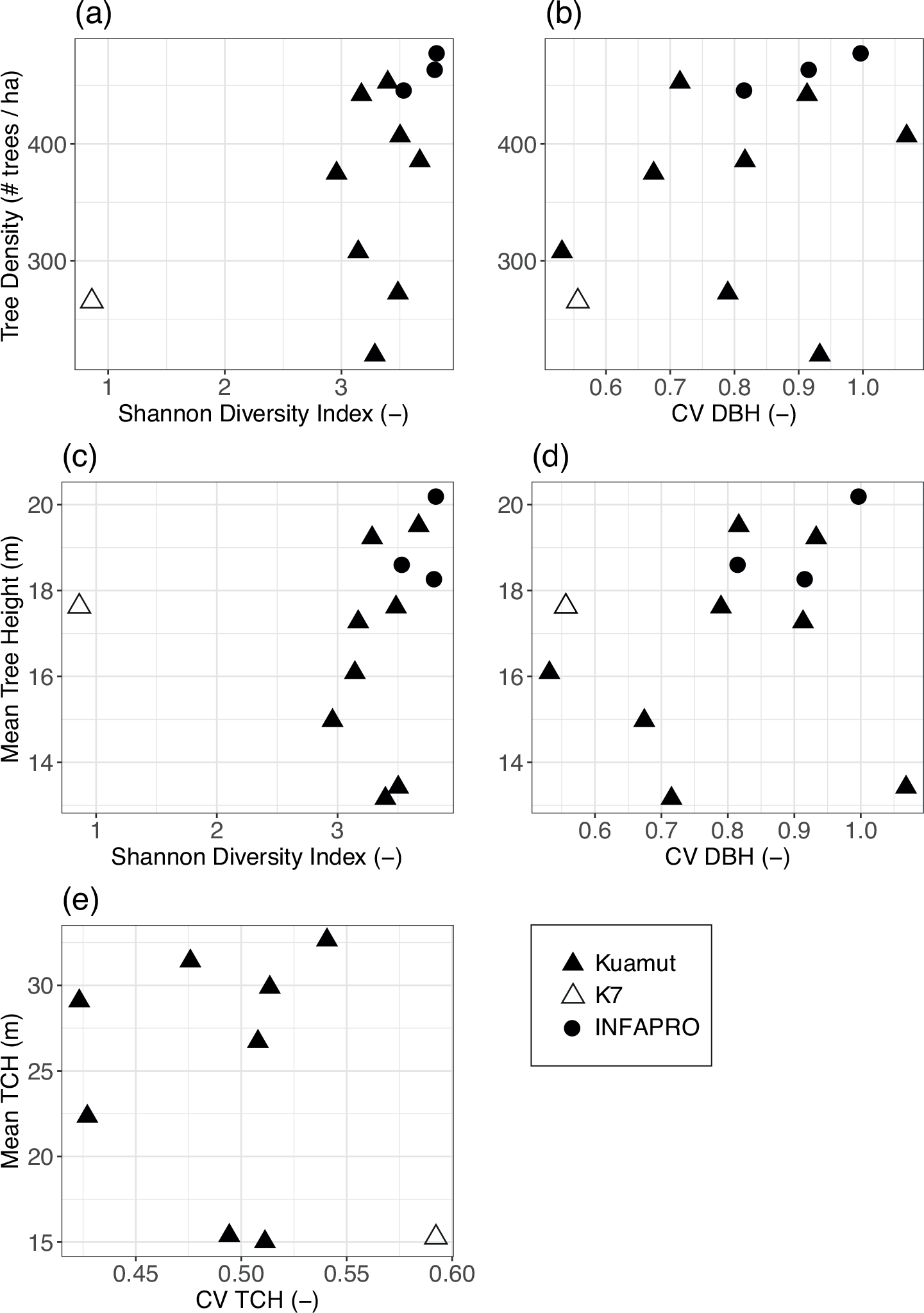
The 12 study plots shown in the forest structural characteristic and diversity space. The top row and middle row show the tree density (a and b) and mean tree height (c and d), respectively, together with the Shannon Diversity Index (a and c) and the coefficient of variation (CV) of Diameter at Breast Height (DBH; b and d) for all plots, while the bottom row shows the LiDAR-derived mean top of canopy height (TCH) and CV of TCH (e) for the plots in the Kuamut area. The plots in Kuamut are indicated by a triangle and the plots at INFAPRO by a circle. Note that LiDAR-data was only available for the Kuamut area. The Shannon Diversity Index was much lower for plot K7 (indicated with an open triangle) than the other plots (see Table 1) but other characteristics did not differ considerably for this plot.

### 4.2. Throughfall

The incoming rainfall for the measured throughfall events varied between 1.4 mm and 126 mm, but events > 100 mm and < 5 mm were uncommon (about 16% of all throughfall measurements). For the majority of throughfall measurements (43-72%, depending on the plot; overall 65%), incoming rainfall was between10 mm and 40 mm (Figure 3). There were fewer throughfall measurements associated with a high rainfall for the plots in the INFAPRO region than for the plots in the Kuamut region. This does not necessarily reflect a climatological difference but is for a large part due to the easier accessibility of the plots in the INFAPRO region, and therefore shorter collection intervals. Often, data collection was not possible directly after a rainfall event in Kuamut (e.g., flooding of the access road), and several rainfall events had occurred by the time the gauges were emptied.

**Figure 3:**
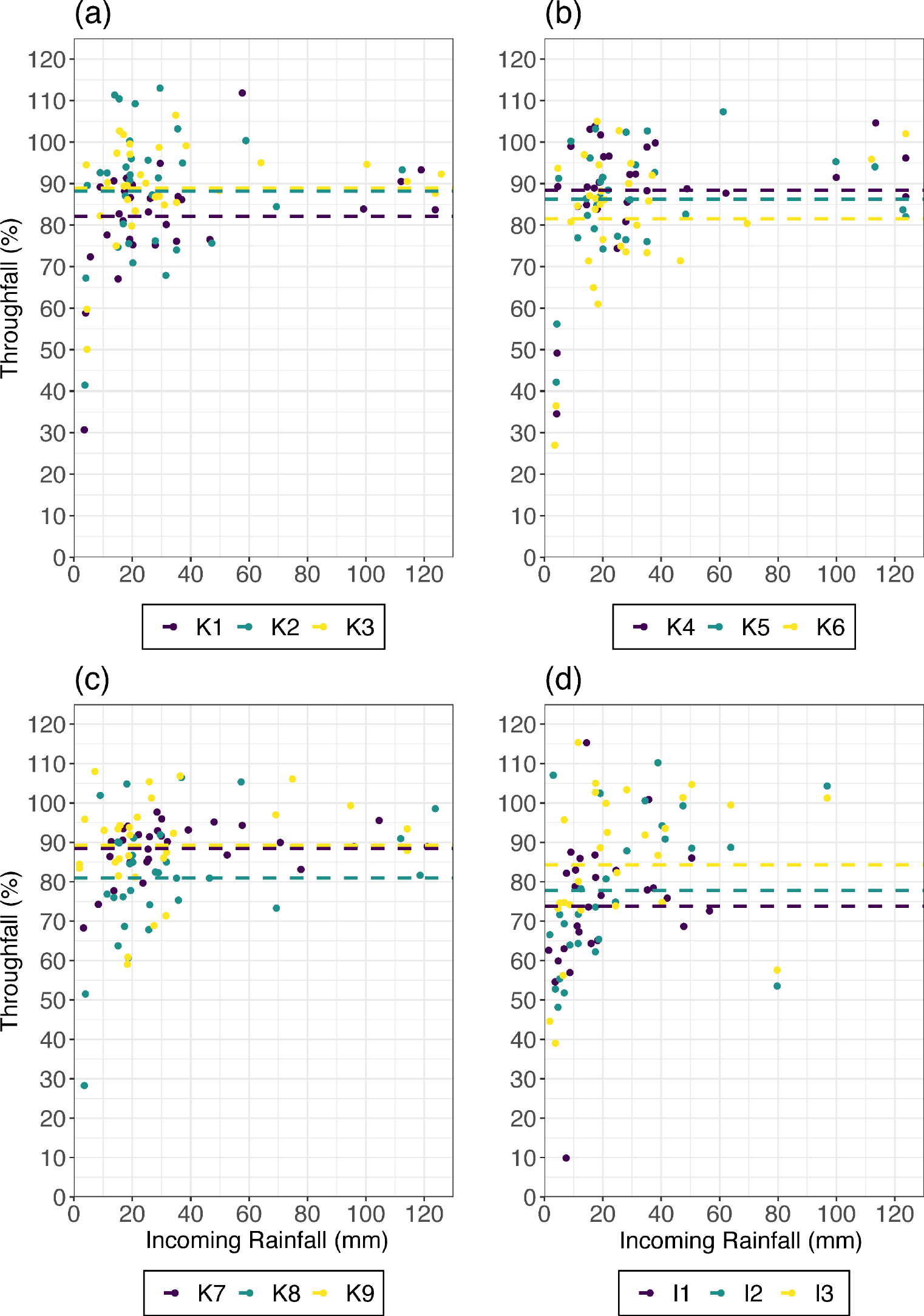
Relation between the plot average throughfall (in % of incoming rainfall) and incoming rainfall (mm) for the 12 study plots (K stands for Kuamut and I stands for INFAPRO, thus K3 is plot 3 in Kuamut). The arithmetic mean throughfall (in %) for each plot is represented by horizontal, dashed line. For reasons of clarity, we show the data of only three plots in each subplot (in numerical order). For the relation between average throughfall and rainfall including the uncertainty in the average throughfall values for each plot, see Figure S5. Table 1 provides detailed information on the plots.

The average throughfall for the measured events varied between 10% to 115% of the incoming rainfall; the interquartile range (i.e., 25^th^ to 75^th^ percentile) of all measurements at all plots ranged from 76% to 94%. There was less throughfall (as a percentage of reference rainfall) for ‘small’ events (< 5 mm) than for periods with more incoming rainfall (Figure 3). The arithmetic mean of the average throughfall (i.e., the mean throughfall) ranged between 74% and 89% (see horizontal dashed lines in Figure 3 and Table 1). The mean throughfall was 79% (± 5%) for the plots in INFAPRO area and 86% (± 3%) for plots in Kuamut (mean for all plots combined: 84%; median: 85%).

### 4.3. Relation between mean throughfall and forest characteristics

#### 4.3.1. Correlations

There was a significant linear and rank correlation between mean throughfall and tree density and basal area (Table 2, and Figures 4 and 5), but no significant relation between mean throughfall and any of the measures related to tree height: ground-derived mean tree height, mean height of the 5% tallest trees, or mean top of canopy height (TCH) (Table 2, and Figures 4 - 6). There was also a significant linear correlation between mean throughfall and the coefficient of variation of the diameter at breast height (CV DBH) and the Shannon Diversity Index (when excluding plot K7; n = 11) (Table 2, Figure 4 and 5). When including plot K7 (n = 12), there was also a significant rank correlation between mean throughfall and Shannon Diversity Index (Table 2). Thus, even though the correlation between mean throughfall and Shannon Diversity Index differs depending on whether plot K7 is included, the overall pattern suggests that mean throughfall and Shannon diversity are related. Mean throughfall was, however, not significantly correlated with the Simpson Diversity Index, coefficient of variation in tree height (CV tree height), and the LiDAR-derived coefficient of variation in top of canopy height (CV TCH)) (Table 2).

**Table 2:** Pearson and Spearman rank correlation coefficients for the relation between the mean throughfall (as percent of incoming rainfall) and structural forest characteristics (fc) and diversity indicators (di), with the corresponding p-values. Significant relations (p < 0.05) are shown in bold font. The results for correlations with the Shannon Diversity Index are reported for two subsets of data (i.e., with and without plot K7; n=12 / n = 11). CV is coefficient of variation. See Table S3 and S4 for the linear model coefficients.

|  |  |  | Pearson | p-value | Spearman | p-value |
| --- | --- | --- | --- | --- | --- | --- |
| ground-derived | forest characteristics | tree density | -0.78 | **0.003** | -0.87 | **< 0.001** |
|  |  | mean tree height | -0.03 | 0.92 | 0.01 | 0.99 |
|  |  | mean tree height (top 5%) | -0.16 | 0.61 | -0.15 | 0.63 |
|  |  | basal area | -0.75 | **0.005** | -0.73 | **0.009** |
|  |  | DBH | -0.10 | 0.76 | -0.14 | 0.67 |
|  | diversity indicators | Simpson Diversity Index | -0.31 | 0.32 | -0.43 | 0.16 |
|  |  | Shannon Diversity Index | -0.45 / -0.60 | 0.14 / **0.05** | -0.60 / -0.54 | **0.04** / 0.09 |
|  |  | CV tree height | -0.33 | 0.29 | -0.34 | 0.29 |
|  |  | CV DBH | -0.58 | **0.05** | -0.53 | 0.08 |
| LiDAR | fc | Top of Canopy Height (TCH) | -0.49 | 0.18 | -0.63 | 0.08 |
|  | di | CV Top of Canopy Height (TCH) | 0.34 | 0.37 | 0.30 | 0.44 |

**Figure 4:**
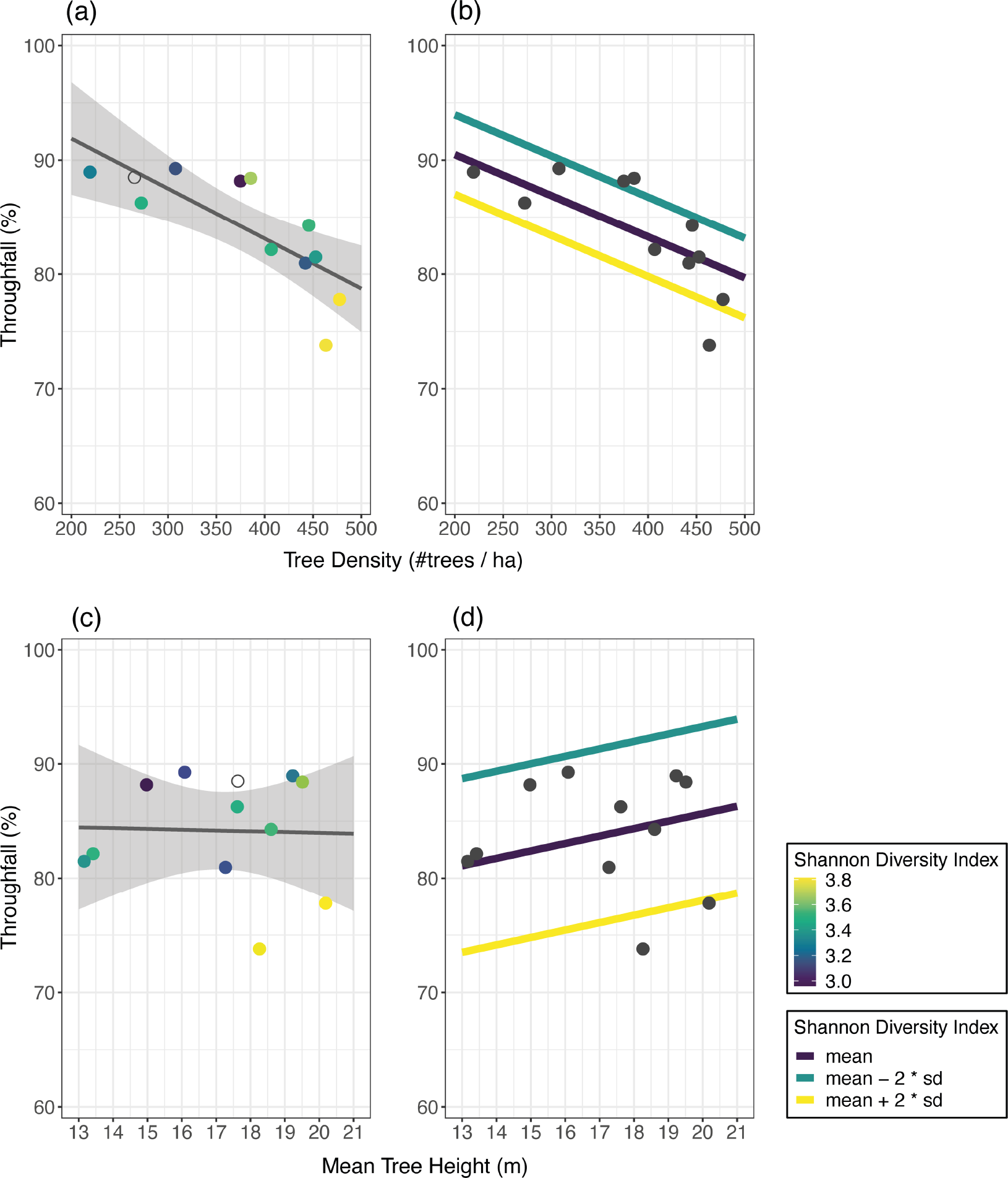
Relation between mean throughfall (in % of incoming rainfall) and tree density (#trees/ha) (a), or the mean tree height (m) (c), and the predicted relationship between mean throughfall and either tree density or mean tree height for different values of the Shannon Diversity Index (b and d). The grey line represents the linear regression line, and the shaded area the 95%- confidence interval. Each dot represents one forest plot and is color-coded according to the Shannon Diversity Index. The colored linear regression lines show the predicted relationships for three scenarios: mean, or mean +/- two times the standard deviation of the Shannon Diversity. Note that one plot (K7) was excluded in the determination of the best-fitting model, and thus not included in subplots b and d. This data point is shown with a grey open circle in subplots a and c.

**Figure 5:**
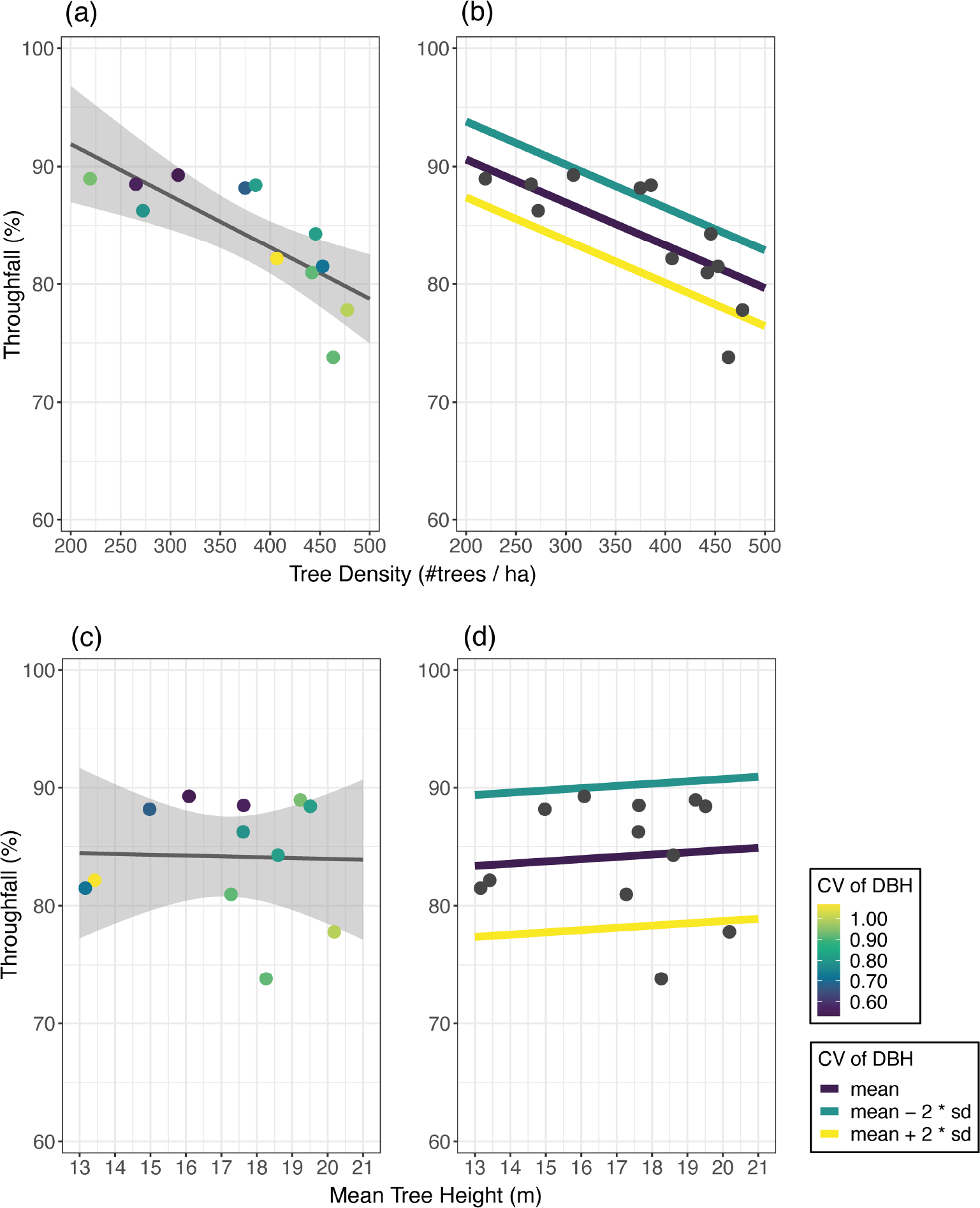
Relation between mean throughfall (in % of incoming rainfall) and tree density (#trees/ha) (a), or the mean tree height (m) (c), and the predicted relationship between mean throughfall and either tree density or mean tree height for different values of the coefficient of variation (CV) of the DBH (b and d). The grey line represents the linear regression line, and the shaded area the 95%-confidence interval. Each dot represents one forest plot and is color-coded according to the CV of DBH. The colored linear regression lines show the predicted relationships for three scenarios: mean, or mean +/- two times the standard deviation of the CV of DBH.

**Figure 6:**
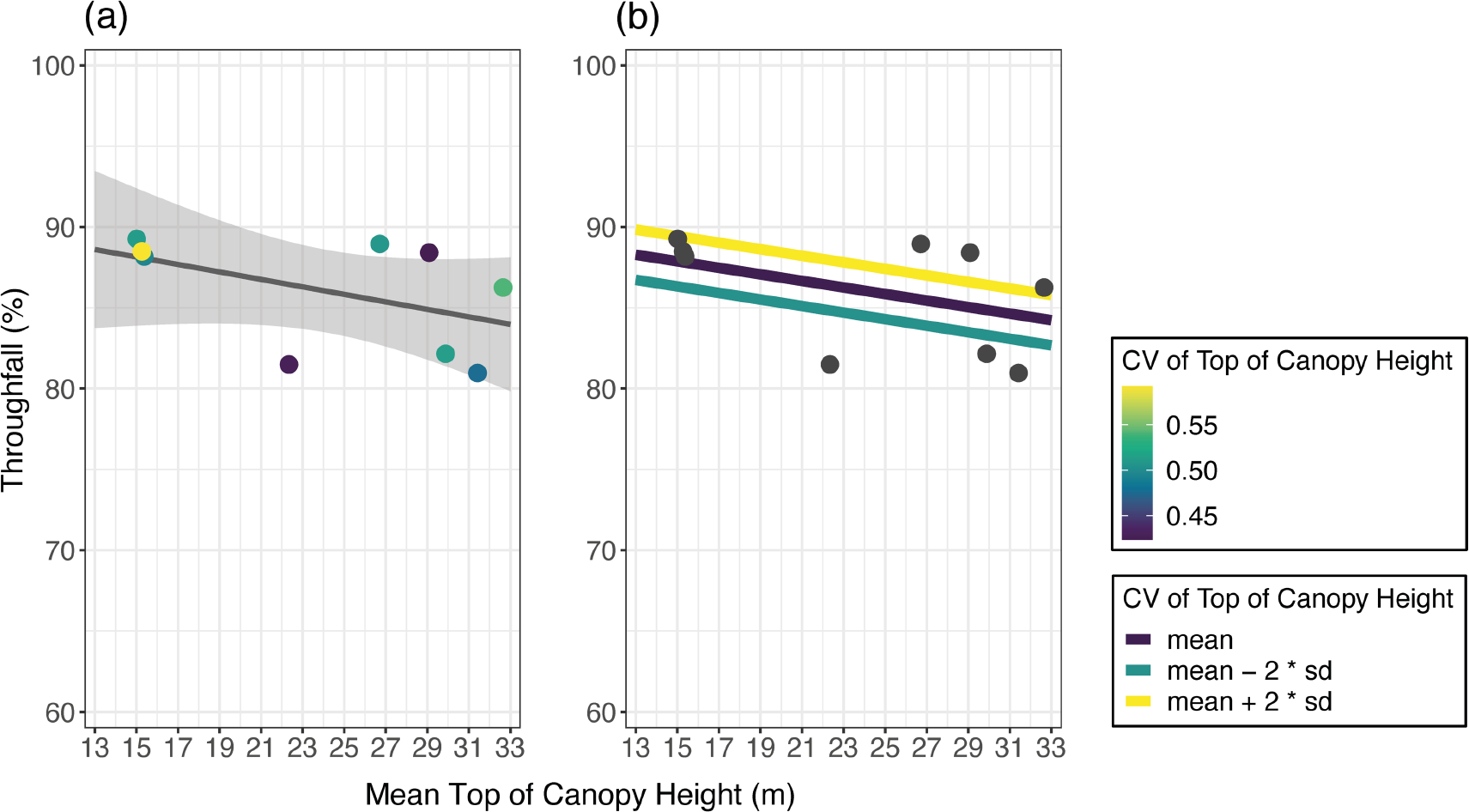
Relation between mean throughfall (in % of incoming rainfall) and the LiDAR-derived mean top of canopy height (m) (a), and the predicted relationship between mean throughfall and top of canopy height for different values of the coefficient of variation (CV) of the top of canopy height (b). The grey line represents the linear regression line, and the shaded area the 95%-confidence interval. Each dot represents one of the nine Kuamut forest plots for which LiDAR-data was available, and is color-coded according to the value of the CV of top of canopy height. The colored linear regressions show the predicted relationship for three scenarios: mean, or mean +/- two times the standard deviation of the CV of top of canopy height.

#### 4.3.2. Linear model for ground-derived forest characteristics

The simple linear model with tree density could explain 56% of the variation in the mean throughfall for the different plots (Figures 4a and 5a; Table S4) but mean tree height explained less than 10% of the variation in mean throughfall (Figure 4c and Figure 5c; Table S4). Consequently, the difference in the AIC values between the null model was only significant (i.e., difference > 2) for the model with tree density (Table 3). This result was consistent for the different subsets of the data (Table 3).

**Table 3:**
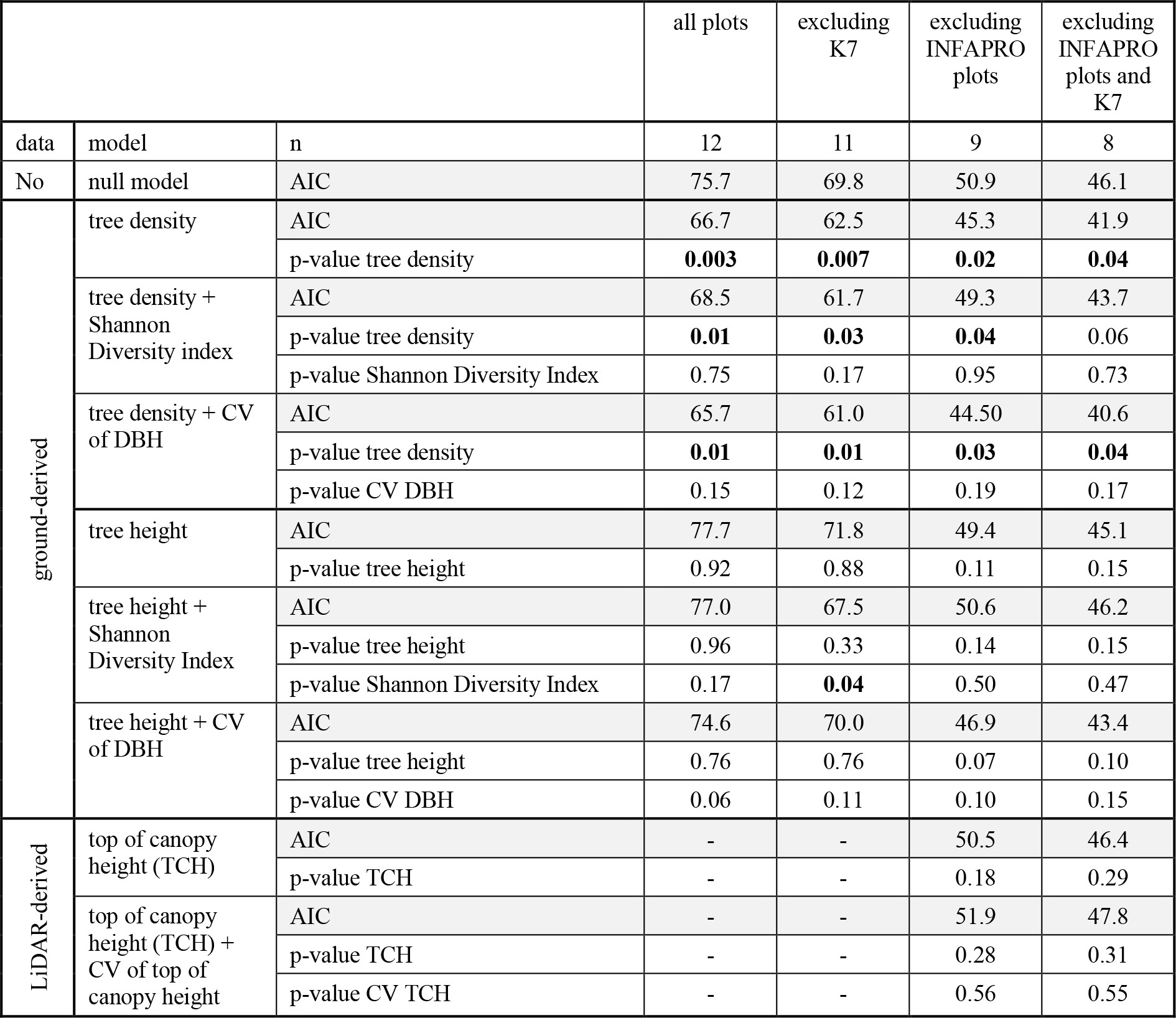
p-values for the linear models explaining the mean throughfall for each plot based on tree density or mean tree height, and when a diversity indicator that reflects forest heterogeneity (Shannon Diversity Index or coefficient of variation (CV) of DBH) is added to the model. Results are also shown for the linear model based on LiDAR-derived mean top of canopy height (TCH) alone and when the CV of TCH is included. There were no significant interaction effects (forest characteristic * diversity measure) for any of the models. Therefore, only the results for the models including the main effects (forest characteristics + diversity indicator) are shown. Significant linear relationships (p < 0.05) are highlighted in bold font. The results for linear models are reported for four subsets of data (in order to obtain comparable AIC values): models with ground-derived data are shown for the full dataset (n=12) and without plot K7 (n = 11); LiDAR-derived data is presented for all plots of Kuamut area (n=9) and without plot K7 (n = 8).

The model including both tree density and Shannon Diversity Index performed better than the null model for the dataset that excludes plot K7 with the high leverage, but only improved the model’s adjusted R-square minimally (up to 0.58), and, as a result, was not significantly better than the simple linear model with tree density alone (Figure 4b; Tables 3 and S4). The linear model for mean throughfall based on the tree density and CV of DBH performed significantly better than the null model, but not significantly better than the model with tree density alone, despite improved model fit (i.e., adjusted R-square increased by 0.06 to 0.62) (Figure 5b, Tables 3 and S4). Furthermore, only tree density had a significant influence on mean throughfall in the multiple linear model that include both the tree density and the CV of DBH (Table 3). There was no interaction effect between either the Shannon Diversity Index or the CV of DBH and the tree density (Table S4).

Adding the Shannon Diversity Index to the linear model with mean tree height improved its performance significantly, so that this model performed better than the null model (excluding K7, Table 3). However, only the Shannon Diversity Index had a significant influence on the mean throughfall in this model and mean tree height did not (Table 3). Furthermore, this multiple linear model could only explain 30% of the observed variation in the mean throughfall for the different plots (Figure 4d; Table S4). The addition of the CV of DBH to the model with mean tree height significantly improved the model performance as well (Table 3). This model could explain 20% of the model’s variance (Figure 5d and Table S4). The CV of DBH had a marginally significant influence on mean throughfall, while mean tree height did not have a significant influence in this model. There was no interaction effect between either the Shannon Diversity Index or the CV of DBH and the tree height (Table S4).

The AIC value was lowest for the multiple linear model with tree density combined with the CV of DBH (Table 3) but this model did not perform better (i.e., difference in AIC values larger than 2) than the simple linear model with tree density alone (Table 3). This result was consistent throughout all the different subsets of data. Further, the models with tree density performed significantly better than the models with mean tree height. Thus, the addition of a diversity measure had little effect on the overall performance of the linear model with tree density and a larger effect on the model with tree density but this model performed not as well as the model with tree density.

#### 4.4.2 Linear model for LiDAR-derived forest characteristics

The linear relation with the mean top of canopy height (TCH) could only explain 13% of the variation in mean throughfall (Figure 6a; Table S4); the difference in AIC values with the null model was not significant (Table 4). The linear model based on the mean TCH and the CV of TCH could only explain 4% of the variation in the mean throughfall for the Kuamut area (Figure 6b; Table S4)). The AIC of this model did not differ significantly to the null-model either, regardless if the AIC was calculated with the data from plot K7 included or not (n=9 or n=8; Table 3). Neither the mean TCH nor the CV of TCH had a significant influence on mean throughfall in this model and there was also no significant interaction between the mean TCH and the CV of TCH (Table S4).

## 5. Discussion

### 5.1. Uncertainty of the throughfall measurements

Despite our efforts to minimize errors (e.g., frequent collection to avoid evaporative losses), we obtained average throughfall values (i.e., for individual plots on specific measurement dates) that were greater than 100%. The mean throughfall values (i.e., averaged over all measurement dates) were, however, less than 100% for all plots. Zimmermann et al. (2013) reported mean throughfall values for two regenerating forest stands that were larger than 100%. Ghimire et al. (2017) attributed this finding to the small number of rain gauges (n = 20) used to measure throughfall. We measured throughfall with 50 rain gauges per plot and moved them three times during the study and therefore assumed that this would be sufficient to adequately capture the spatial variability in throughfall. To test this assumption, we calculated the number of gauges required to obtain a margin of error smaller than ±10% with a 90% confidence following the method of Holwerda et al. (2006). For a standard of deviation of 36% (the mean of the standard deviation for all plots and collection dates), the minimum sample size is 63. For 10% of the measurements the standard deviation was less than 24% and 50 gauges would have been sufficient. Thus, despite the large number of gauges used in this study, this may not have been sufficient to determine the average throughfall with 90% certainty on some of the measurement dates.

As Ghimire et al. (2017) point out, a high gauge density also increases the probability of measuring throughfall directly under so-called dripping points (i.e., funneling of rainfall in the canopy), which generally leads to a higher average throughfall (see also Holwerda et al. 2006; Lloyd et al. 1988). The measurements with a higher average throughfall than incoming precipitation were not all from the same period (i.e., before moving the gauges) and were also not caused by one or two gauges with extremely high throughfall. We, therefore conclude that they were not caused by extremely high throughfall in gauges below drip points. Rather, we assume that the very high throughfall values are caused by errors in the measurements of the incoming rainfall. We tried to put the gauges at least 20 m (or ∼1.5 times the tree height) away from the forest edge but open locations were strongly limited. Strong winds may have caused an under-catch of the reference rainfall gauges for some of the measurement dates. Alternatively, the distance between the reference rainfall gauges and the forest plots (max 536 m) may have been too large or the inverse distance weighting method was inappropriate. However, the average throughfall was more than 100% for 17% of the measurements at plots with nearby rain gauges for which no inverse distance weighting was used (varying between 0 and 32%) vs. 13% of the measurements for plots for which inverse distance weighting was used (varying between 3 and 27%). This suggests that the distance weighting method was not the main source of error. Based on the somewhat similar fraction of measurements for which throughfall was larger than incoming precipitation for the different plots, we assume that these errors did not lead to a large systematic difference in the mean throughfall for the different plots.

Alternatively, fog interception may have contributed to the high throughfall. Bruijnzeel et al. (2011) estimated a mean annual fog input between 50 – 100 mm (i.e., 2.5 – 5% of total precipitation (rainfall plus fog)) for the Kuamut and INFAPRO region. We did not account for fog input, but it is likely that this contributed unequally to the rain gauges in the forest and in the open areas and would lead to systematic differences between the forest plots. However, we assume that this effect is relatively small due to the overall small contribution of fog to total precipitation. Furthermore, the percentage of measurements with an average throughfall between 100% and 120% of precipitation was not related to tree density, tree height, the Shannon Diversity Index of the CV of DBH (p = 0.20, 0.37, 0.94, and 0.96 respectively).

Ten measurement dates were excluded from the calculation of the mean throughfall because the average throughfall was >120% of the incoming rainfall. This represents 2.7% of all measurements at the plots. The very large differences between mean throughfall and incoming rainfall are also unlikely to be caused by fog drip. We, therefore, considered these measurements to be influenced by measurement errors, most likely in the open rainfall. They were unlikely to be caused by drip points because they were not caused by a high throughfall in a few gauges.

### 5.2. Comparison of mean throughfall for the plots with other studies

The overall mean throughfall for all plots (84 ± 5%) was slightly higher than the value (81%) reported by Sinun et al. (1992) for throughfall in a primary lowland dipterocarp rain forest at the East Ridge Area in Danum Valley Conservation Area (DVCA) (Palum Tambun catchment; described by Walsh et al. (2011)). The mean throughfall for this study is also similar to the average throughfall for a lowland dipterocarp rain forest that was selectively logged with different levels of disturbance (but not yet classified as cleared areas) near DVCA: 80 ± 7% to 84± 7% (Chappell et al. 2001). Chappell et al. (2001) also report a substantially higher (91 ± 7%) mean throughfall for an undisturbed (virgin) protected forest than Sinun et al. (1992), although their measurements were influenced by the 1997 – 1998 El Niño during which there was substantial leaf loss. Thus, despite the relatively large number of measurement points for which average throughfall was larger than the reference precipitation (see section 5.1), the mean throughfall values for the plots seem reasonable.

The plots studied here were less dense (mean 376 trees per ha; range 219-478 trees per ha) and diverse (mean Shannon diversity of 3.22, range: 0.86-3.81) than reported by Hayward et al. (2021) and in previous throughfall studies. For example, Hayward et al. (2021) reported a tree density of 460 trees per ha, and a mean Shannon Diversity Index of 4.81 for logged-over forest plots at INFAPRO region. Old growth forest plots at Danum Valley Conservation Area had a tree density of 590 trees per ha with a mean Shannon Diversity Index of 4.56 (Hayward et al. 2021). The reported tree height, density and Shannon Diversity Index for secondary forests outside Borneo where throughfall has studied varied between 6.6-19 m, 695-936 individuals per ha, and 16-35, respectively (Dietz et al. 2006; Geißler et al. 2013; Ghimire et al. 2017; Ponette-González et al. 2010). Note, however, that the calculations of these values differ (e.g., plot size, minimum DBH to be included in the sample size) and, therefore, need to be compared to our values with caution. Moreover, forests might differ in their management and restoration history, and are thus, again only comparable to a limited extent.

### 5.3. Variation in mean throughfall across the regenerating logged-over forest landscape

The variation in mean throughfall as a percentage of incoming precipitation for our study plots (range: 74-89%) suggests that throughfall is highly variable across the regenerating tropical forest landscape. Mean throughfall was lower for the plots in the INFAPRO area than the Kuamut area (ANOVA-test, p=0.02, F-statistics: 8.1 on 1 and 10 DF), suggesting that time since disturbance affects mean throughfall. Plots in the INFAPRO area had a longer recovery time (30 - 50 years) than in Kuamut (10 - 15 years) and underwent passive, as well as active ecosystem restoration (i.e., enrichment planting, climber cutting). The mean throughfall values for the plots in INFAPRO (79±5%) approach the values reported for primary forest by Sinu et al. (1992). The observed difference in mean throughfall does not depend on the frequency of the measurements and holds when the mean throughfall is calculated only for collection dates with an incoming reference rainfall < 100 mm (i.e., when using the same range of incoming precipitation for both areas; Figure 3). This change in mean throughfall with plot age for the studied secondary lowland dipterocarp forests plots is slower than for the semi-deciduous lowland forests in Panama, where throughfall values approached those of mature forests within ten years (Zimmermann et al., 2013). Our study design does not allow the separation of the effects of age or/and restoration management (i.e., active restoration vs. natural regeneration) because of (for that purpose) unbalanced experimental designing and too small sample size.

Instead, we sought the relationships between forest characteristics and mean throughfall on th island of Borneo and highlighted the important role of tree density (Figure 4 and Figure 5). In our study, basal area was correlated with tree density (Figure 1 and Table S2) and also significantly correlated with throughfall (Table 2). This result is contrary to the results for studies for secondary forest sites in lowland tropical rainforests in Panama by Zimmermann et al. (2013) and a study in a lower montane rainforest in Central Sulawesi (Indonesia) by Dietz et al. (2006), who did not find such a relation between mean throughfall and basal area. Unlike Dietz et al. (2006) and Ponette-González et al. (2010), we did not find a significant relationship between mean throughfall and mean tree height, and the mean minimum tree height, respectively. Nevertheless, Zimmermann et al. (2013) and Ponette-González et al. (2010) found a significant relationship between throughfall and basal area for a lowland forest in Panama a montane forest in Mexico, but Dietz et al. (2006) did not observe such a relation with basal area. Dietz et al. (2006) did report a significant correlation between mean DBH on throughfall, but did not report on whether DBH was correlated with any other forest parameters. For the plots studied by Zimmermann et al. (2013), tree density and tree basal area were only correlated for stems with a DBH <5 cm (which were not included in our plot surveys), but not for large trees (i.e., DBH >5 cm). Ponette-González et al. (2010) did not report on the relationship between throughfall and tree density.

### 5.4. Effect of heterogeneity of diversity on mean throughfall

There was a significant correlation between the Shannon Diversity Index and mean throughfall (Table 2), and a slight improvement in performance (i.e., adjusted R-square) of the linear model with mean tree height when including the Shannon Diversity Index. However, this model did not perform significantly better (i.e., difference in AIC < 2; Table 3) than the overall best-performing models (i.e., model with tree density alone or in combination with the CV of DBH). Zimmermann et al. (2013) is (to our knowledge) the only other study that investigated influence of diversity on throughfall, but they did not find a correlation between Shannon Diversity Index and throughfall for their forest plots in Panama. As they conducted their study in differently sized plots, it is likely that their Shannon Diversity Index values were not fully comparable among themselves (because larger sampling area have a higher likelihood of encountering rare species) (cf. Budka et al., 2019). Geißler et al. (2013) showed a significant effect of tree diversity (i.e., rarefied tree species richness and functional diversity) on throughfall kinetic energy in mixed evergreen broad-leaved subtropical forest consisting of different successional stages (forest described by X. Liu et al. 2018) in the Zhejiang Province in China.

Mean throughfall was also related to the variation in DBH, and inclusion of this diversity indicator indeed led to an increase in model performance. However, this increase was only statistically significant for the model with mean tree height. We note that the simple linear model with mean tree height alone had very low explanatory power.

The importance of explanatory variables related to the basal area (and in our dataset therefore also the CV of DBH due to the high correlation between these two variables, see Figure 1) for throughfall models was already highlighted by Zimmermann et al. (2013). Their best fitting model for throughfall was based on the ratio of basal area of small (DBH <5 cm and >1 cm) and large (DBH >5 cm) trees. As our structural forest data only included stems with a DB >10 cm, we were not able to calculate this ratio. The influence of the CV of DBH on throughfall supports the claim of Zimmermann et al. (2013) that variables related to basal area should be included in future throughfall-modelling efforts. The comparison of our results with those of previous studies (i.e., (Dietz et al. 2006; Ponette-González et al. 2010; Zimmermann et al. 2013) suggests that forest parameters related to tree density, basal area, or DBH are particularly relevant to understanding the variation in mean throughfall across the regenerating degraded forest landscape. In view of the limited explanatory value of CV of DBH and tree species diversity, and the large effort required for species identification, we regard the additional value of including diversity indicators in linear models to estimate throughfall is minor. Based on our results, it seems that tree density alone is sufficient to estimate the regional variation in mean throughfall.

This study focused on the effect of plot characteristics on the overall, arithmetic mean throughfall (cf. Chappell et al. 2001; Zimmermann et al. 2013; Ghimire et al. 2017). The effect of (structural) forest diversity may be greater for smaller rainfall events because the effect of vegetation density is larger for the events that do not fully saturate the canopy. Unfortunately, we could not test this hypothesis with our data because only 9% of all measurements (33 in total) were for small (< 5mm) events. Nevertheless, distinguishing between small and large rainfall events may allow for a better insight in forest-rainfall interactions, particularly with regards to canopy wetting and saturation processes during small rainfall events. A future focus on small rainfall events would therefore be instructive.

### 5.5. Use of LiDAR-derived Top of Canopy (TCH) data to estimate throughfall across the landscape

LiDAR-derived mean top of canopy height (TCH) could not explain the variation in mean throughfall, even including the variability in the top of canopy height (i.e., CV of TCH). Several previous studies have suggested that LiDAR-derived canopy data can be used to estimat throughfall. However, most of these studies focused on LAI (e.g., Teske and Thistle 2004), or canopy closure (e.g., Roth et al. 2007) instead of the top of canopy height. For example, Teske and Thistle (2004) mapped LAI (obtained, among other sources, by LiDAR data) for forests of North America and demonstrated the utility of LiDAR data to estimate interception at a landscape scale. We did not have LAI data, and focused on the top of canopy height assuming this would be a proxy for time since regeneration or degree of degradation.

A recent study on snow interception in a Swiss mountain forest found relationships between interception and indices that describe canopy gaps (i.e., “mean distance to canopy”, and “total gap area”) (Moeser et al. 2015). These measures of canopy gaps could be very relevant in estimating throughfall in diverse tropical forests as well. However, the studied forest plots were too dense and too small to detect gaps. Furthermore, the TCH map was only two-dimensional and had a (for this sort of application) comparably large spatial scale (i.e., 1 m). Nevertheless, three-dimensional LiDAR data at very high resolution may be useful to determine the effects of fine scale gap structure on throughfall. This potential was already highlighted by Roth et al. (2007). Other studies have shown the usefulness of this type of data for three-dimensional models of microclimate (Broadbent et al. 2014) and changes in forest structure due to biological invasions (Asner et al. 2008).

## 6. Conclusions

Throughfall measurements at twelve logged-over forest plots in Borneo show that average throughfall varies significantly across the landscape and that vegetation regrowth will thus affect the water balance for this area. Mean throughfall was closely related to tree density, basal area, Shannon diversity, and the variation in DBH, but some of these forest variables were also highly correlated with each other. Mean throughfall was not significantly related to mean tree height (field measurements) or the mean top of canopy height (LiDAR-derived measurements). The linear regression between forest structural characteristics could in some cases be improve by adding a measure of diversity but it did not improve the model beyond the simple linear regression with tree density. These results highlight the importance of tree density and suggest a limited effect of diversity on throughfall.

## Contribution of authors

This project was conceptualized by NK, IvM, and CDP. The throughfall data were collected by NK, data on the forest stands were collected by NK and CDP, and the LiDAR data were provided by GPA. The manuscript was written by NK, IvM, CDP, and JG, and supported by EG, HT, GPA. There is no conflict of interest.

## Acknowledgements

We thank the team of research assistants from Permian Global and the South-East Asian Royal Research Partnership (SEARRP) for their tremendous help in setting up the rain gauges, and collecting the throughfall and rainfall data: terima kasih, Abang Philip, Joulu, Gapik, Unjai, Weldy, France, dan Eli. We further thank the Sabah Forestry Department for their efforts in mending the road, lending us their vehicle, and allowing us to camp in one of their checkpoint houses. We are, furthermore, grateful for the help with coding and statistical advice by Navjot Sidhu and Vasileios Tsakalos.

The project was funded by ETH Zurich (Switzerland) grant ‘ETH-36 16-2, FORESTeR’ to CDP, which provided funding for NK’s PhD studentship. The research was conducted under Sabah Biodiversity Council Access Licence (Licence Ref.No.: JKM/MBS.1000-2/2 JLD. (45)) to CDP. Global Airborne Observatory mapped and data processing was supported by the UN Development Programme GEF, Avatar Alliance Foundation, Roundtable on Sustainable Palm Oil, Worldwide Fund for Nature, Morgan Family Foundation, and the Rainforest Trust. D.A.

## Supporting Information

Supplementary material 1 provides addition information on the forest characteristics analyzed in this study and their correlation.

Supplementary material 2 provides more information on the design of the rainfall gauges and the evaluation of evaporation from the gauges.

Supplementary material 3 provides additional analyses of the throughfall data. Supplementary material 4 provides additional information on the best fitting linear models.

