## Supplementary material 1, 2, 3, and 4 for "Does forest heterogeneity affect mean throughfall for regenerating secondary forests on Borneo?"

**Supplementary material 1: Forest characteristics analysed in this study**

Table S1: List of all forest characteristics and diversity indicators (based on field data and LiDAR-data) used in this study.

|  | | **Abbreviation** | **Description** | **Units** |
| --- | --- | --- | --- | --- |
| ground-derived | forest characteristic | tree density | Number of trees per plot multiplied by 1/*plot_size* in order to obtain density per hectare | # trees / ha |
|  |  | tree height | Mean tree height per plot | m |
|  |  | tree height (top 5%) | Mean tree height of the 5% tallest trees | m |
|  |  | basal area | Sum of the surface area of all stems at 1.30 m height | m^2^ |
|  |  | DBH | Mean DBH per plot | cm |
|  | diversity indicator | Simpson Diversity Index | Simpon Diversity Index calculated with the *diversity()*-function of the R package *vegan* (Oksanen *et al.*, 2020). *simp* was calculated on the species level using data for all identified trees. Trees that were only identified at the genus-level were treated as one species. Unidentified species and lianas were grouped together and treated as one species, as well. | - |
|  |  | Shannon Diversity Index | Shannon Diversity Index calculated with the *diversity()*-function of the R package *vegan* (Oksanen *et al.*, 2020). *shan* was calculated on the species level using data for all identified trees,. Trees that were only identified at the genus-level were treated as one species. Unidentified species and lianas were grouped together and treated as one species, as well. | - |
|  |  | CV tree height | Coefficient of variation of tree height; calculated as standard deviation of tree height divided by the mean tree height | - |
|  |  | CV DBH | Coefficient of variation of tree height; calculated as standard deviation of DBH divided by the mean tree height | - |
| LiDAR-derived | forest characteristic | TCH | Mean top of canopy height (TCH) per plot | m |
|  | diversity indicator | CV TCH | Coefficient of variation of TCH; calculated as standard deviation of TCH divided by the mean TCH | - |

Table S2: p-values of correlation tests (following Pearson distribution (light-grey shaded) and Spearman rank distribution (unshaded)) between all ground-derived structural forest characteristics (fc) (tree density, tree height, tree height (top 5%), basal area, DBH) and diversity indicators (di) (Simpson Diversity Index, Shannon Diversity Index, CV tree height, CV DBH), as well as the LiDAR-derived top of canopy height (TCH) and CV of TCH). Significant p-values (i.e., p < 0.05) are highlighted in bold font). Note that the results with the Shannon Diversity Index are reported for two subsets of data, including and excluding plot K7 (n=12 / n = 11). For the correlation coefficients, see Figure 1 and S1.

|  | | | tree density | tree height | tree height (top 5%) | basal area | DBH | Simpson Diversity Index | Shannon Diversity Index | CV tree height | CV DBH | TCH | CV TCH |
| --- | --- | --- | --- | --- | --- | --- | --- | --- | --- | --- | --- | --- | --- |
| ground-derived | forest characteristics | tree density |  | 0.75 | 0.72 | **0.01** | 0.60 | 0.18 | 0.10 / 0.19 | 0.60 | 0.19 | 0.63 | **0.04** |
|  |  | tree height | 0.85 |  | **< 0.01** | 0.28 | **0.01** | 0.96 | 0.76 / 0.13 | 0.80 | 0.53 | 0.54 | 0.74 |
|  |  | tree height  (top 5%) | 0.70 | **< 0.01** |  | 0.14 | **< 0.01** | 0.31 | 0.19 / 0.19 | **0.04** | 0.06 | 0.19 | 0.92 |
|  |  | basal area | **0.01** | 0.14 | **0.03** |  | 0.08 | 0.14 | **0.03 /**  **< 0.01** | **0.03** | **< 0.01** | **0.03** | 0.17 |
|  |  | DBH | 0.70 | **< 0.01** | **< 0.01** | **0.03** |  | 0.62 | 0.37 / **0.04** | 0.07 | 0.05 | 0.20 | 0.90 |
|  | diversity indicators | Simpson Diversity Index | 0.09 | 0.12 | **0.05** | **0.01** | **0.02** |  | **< 0.01 / < 0.01** | 0.08 | 0.08 | 0.17 | **0.05** |
|  |  | Shannon Diversity Index | **0.02** / **0.05** | 0.08 / 0.06 | **0.02** / 0.06 | **< 0.01 / < 0.01** | **0.01 / 0.02** | **< 0.01 / < 0.01** |  | 0.06 / 0.31 | 0.06 / 0.08 | 0.09 / 0.07 | **0.04 /** 0.50 |
|  |  | CV tree height | 0.44 | 0.29 | **0.01** | **0.01** | **0.04** | 0.09 | **0.05** / 0.15 |  | **< 0.01** | 0.08 | 0.59 |
|  |  | CV DBH | 0.21 | 0.24 | **0.02** | **< 0.01** | **0.02** | 0.06 | **0.02** / 0.06 | **< 0.01** |  | 0.08 | 0.52 |
| LiDAR | FC | TCH | 0.52 | 0.74 | 0.10 | **0.04** | 0.21 | 0.18 | 0.06 / 0.15 | 0.11 | **0.03** |  | 0.48 |
|  | DI | CV TCH | **0.09** | 0.95 | 0.68 | 0.27 | 0.88 | 0.52 | 0.39 / 0.93 | 0.44 | 0.64 | 0.84 |  |

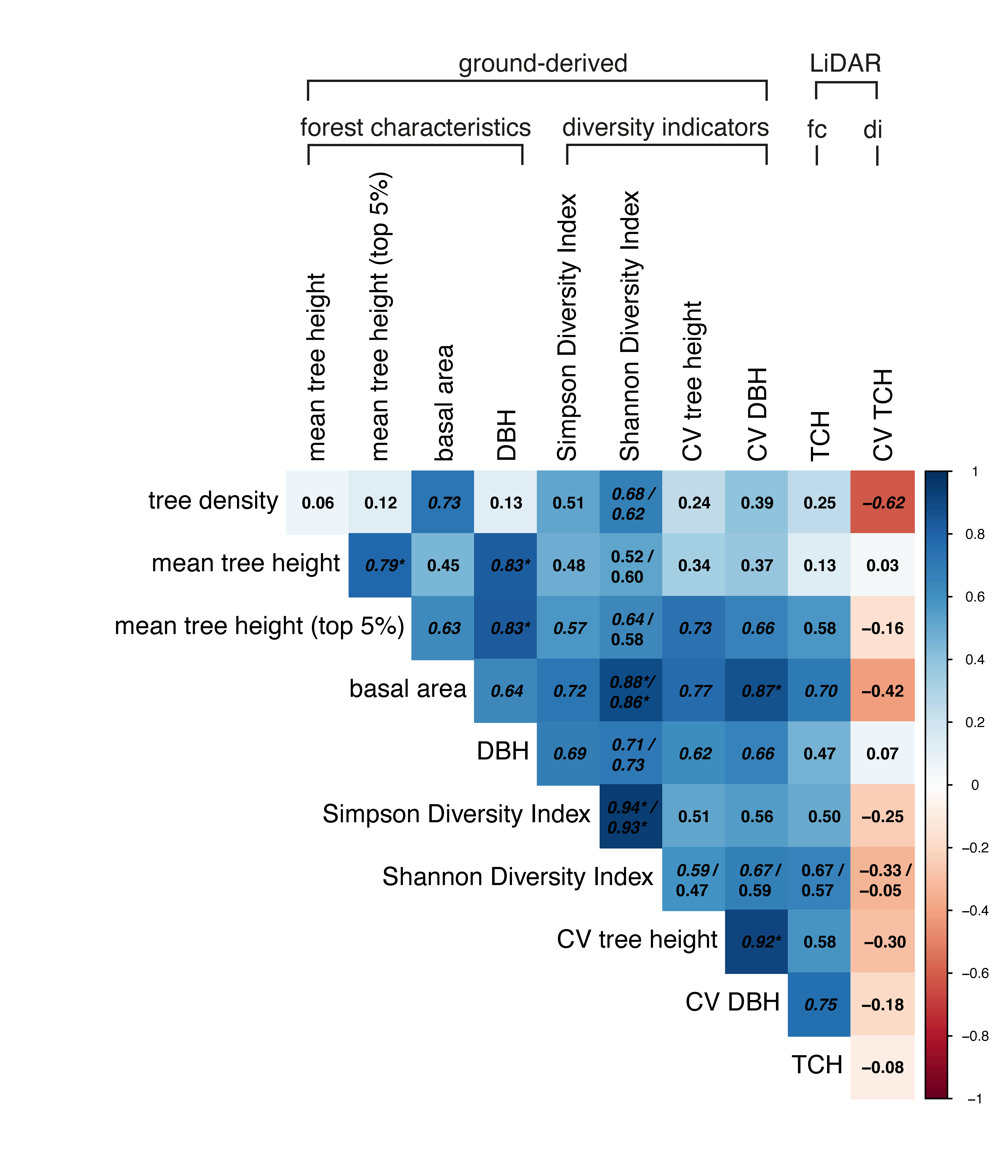

Figure S1: Spearman correlation coefficients between all ground-derived structural forest characteristics (fc) (tree density, tree height, tree height (top 5%), basal area, DBH) and diversity indicators (di) (Simpson Diversity Index, Shannon Diversity Index, CV of tree height, and CV of DBH), as well as the LiDAR-derived Top of Canopy Height (TCH) and CV of TCH. Significant (p<0.05) correlations are italicized and correlations with p<0.01 are additionally marked with a * (see Table S2 for p-values). Correlation values for Shannon Diversity Index are reported for two subsets of data (n=12 / n = 11). Note that the color of the tiles refers to the correlation tests with the full dataset (n=12).

**Supplementary material 2: Design and evaluation of the rainfall gauges**

For the rain gauges, we used 1.5 L plastic bottles. We cut the upper part of the bottle and placed it upside down in the bottle, so that it functioned as a funnel (cf. Hendriks 2010; Davids et al. 2019). We replaced the lid with pantyhose in order to avoid debris and litter falling into the bottle (Figure S3). All bottles had the same diameter and were fixed with wire on wooden poles at 1 m above the ground to avoid ground-splash (cf. Ghimire et al. 2017).

Although the initial wetting of the plastic bottle and the pantyhose causes some underestimation of the amount of throughfall (particularly for small (<5 mm) rainfall events), we assume that this effect is small because the reference rainfall measurements would be affected by the same losses. Furthermore, only 33 out of the analyzed events (n = 361) were < 5 mm.

Because the gauges were emptied daily, and occasionally up to 72 hours after the event, the measurements can also be influenced by evaporation from the gauges. In particular, the measurements of the reference rainfall may have been affected by evaporation as they were more exposed to the sun. This would lead to an underestimation of the incoming rainfall, and therefore overestimation of the average throughfall as a percentage of rainfall. We, therefore, filled four rain gauges with water and compared evaporation losses in the open field and the forest. The evaporation rate (average of ~ 0.07 mm/d) only diverged after the third day, and therefore, only measurements taken within 72 hours after the rainfall event are included in the analyses.

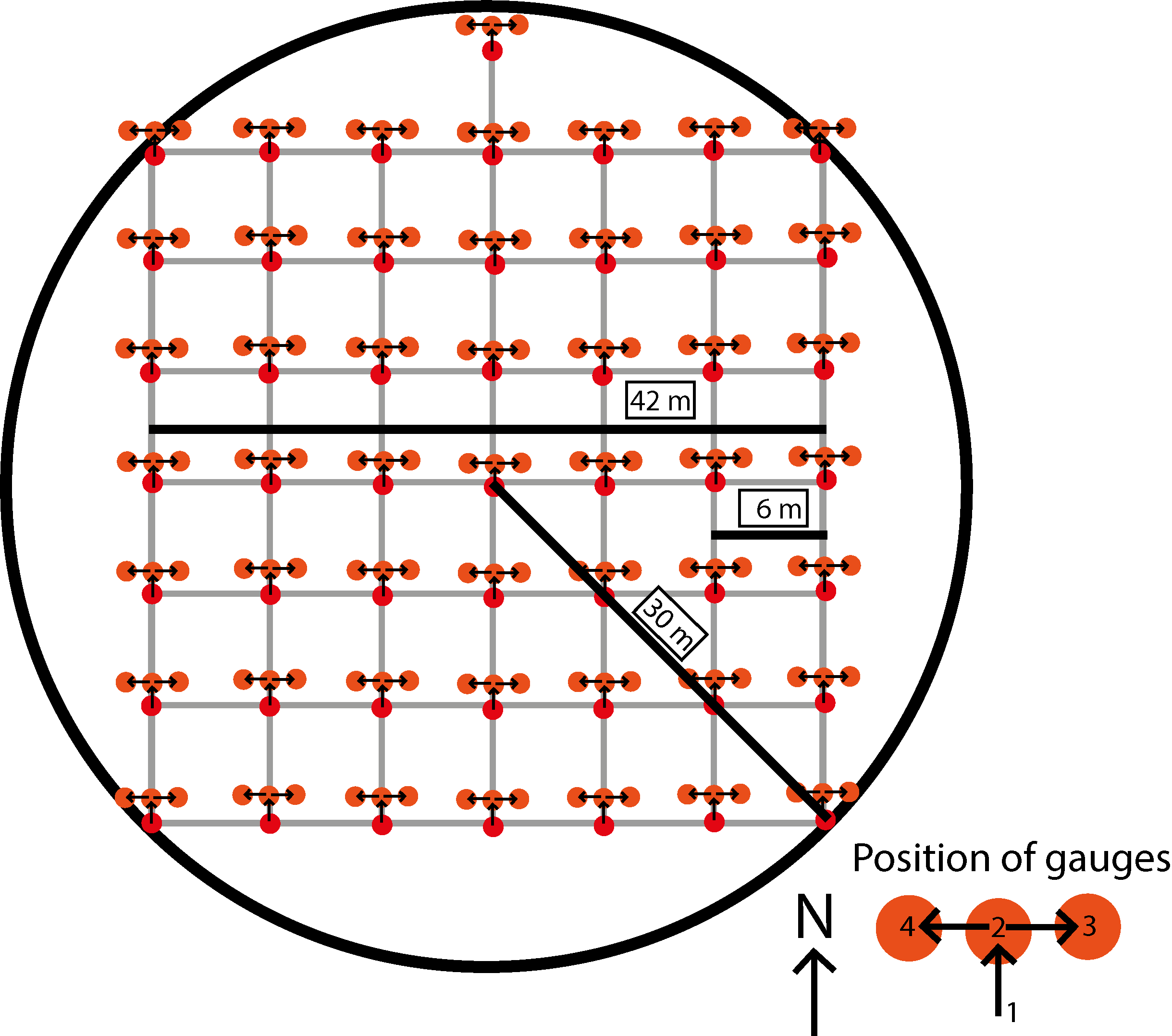

Figure S2: Set up of the rain gauges (red dots) on the regular 7x7 grid (grey lines) within the forest plot (black circle). The thick, black lines provide information about the distances: the radius of the forest plots was 30m, the total width/length of the regular grid was 42 m, and the distance between individual rain gauges was 6 m. The rain gauges were moved three times during the study period (after 40 days 1 m to the north, after 80 days 1 m to the east, and after 120 days 2 m to the west (1 m to the west from second position)), as indicated by the orange-colored circles and the legend. Please note that even though the two rain gauges in the top right corner and top left corner of the appear to be moved outside the borders of the plot on position three, respectively four, this was not the case. Instead of moving them one meter, we just moved them about 10 – 20 cm to stay within the borders of the plot.

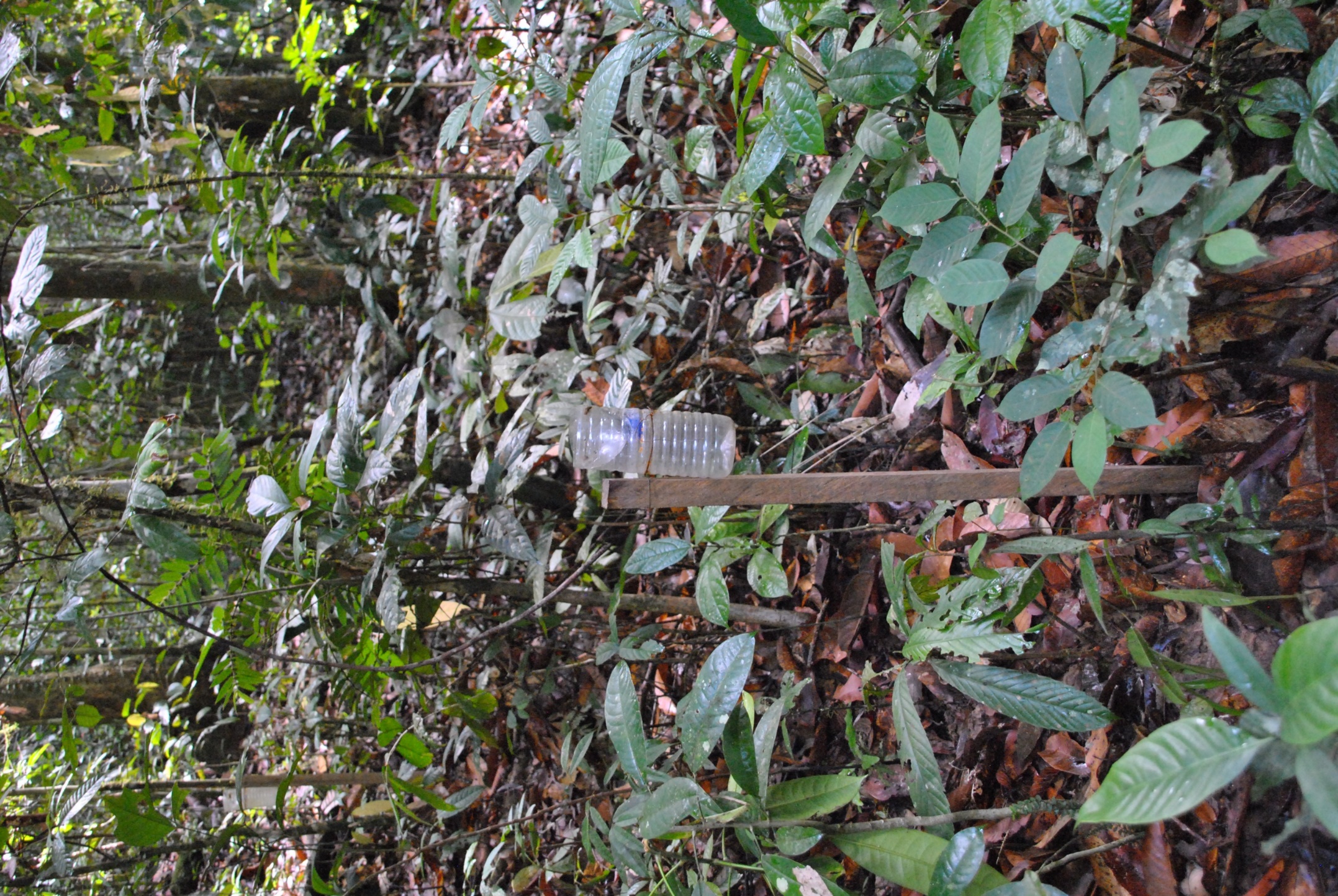

Figure S3: Photo of the rain gauge consisting of a cut plastic bottle, with the upper part functioning as funnel placed inside the bottle, and fixed with wire at 1 m above ground on a wooden pole.

**Supplementary material 3: Throughfall outlier detection**

We determined the reliability of the average throughfall measurements for individual measurement dates by plotting the average throughfall (in mm) against reference rainfall (in mm) (Figure S4). Data points that clearly diverged from the expected linear regression line were re-examined for missing data or typos. It is not unusual to obtain throughfall values exceeding the size of the reference rainfall values (see reasoning in main text); yet, we applied a threshold of TF = 120%, and considered measurements above this value (n = 10) to be erroneous, and consequently excluded from the further analysis. This had only a small effect on the calculated mean throughfall and changed the linear model between throughfall (in mm) and reference rainfall (in mm) only slightly (model for all data: R^2^ = 0.97, intercept: -0.89, slope: 0.92, p < 0.001; model after exclusion of 10 datapoints with mean throughfall values above 120%:(R^2^ = 0.98, intercept: -1.14, slope: 0.92, p < 0.001

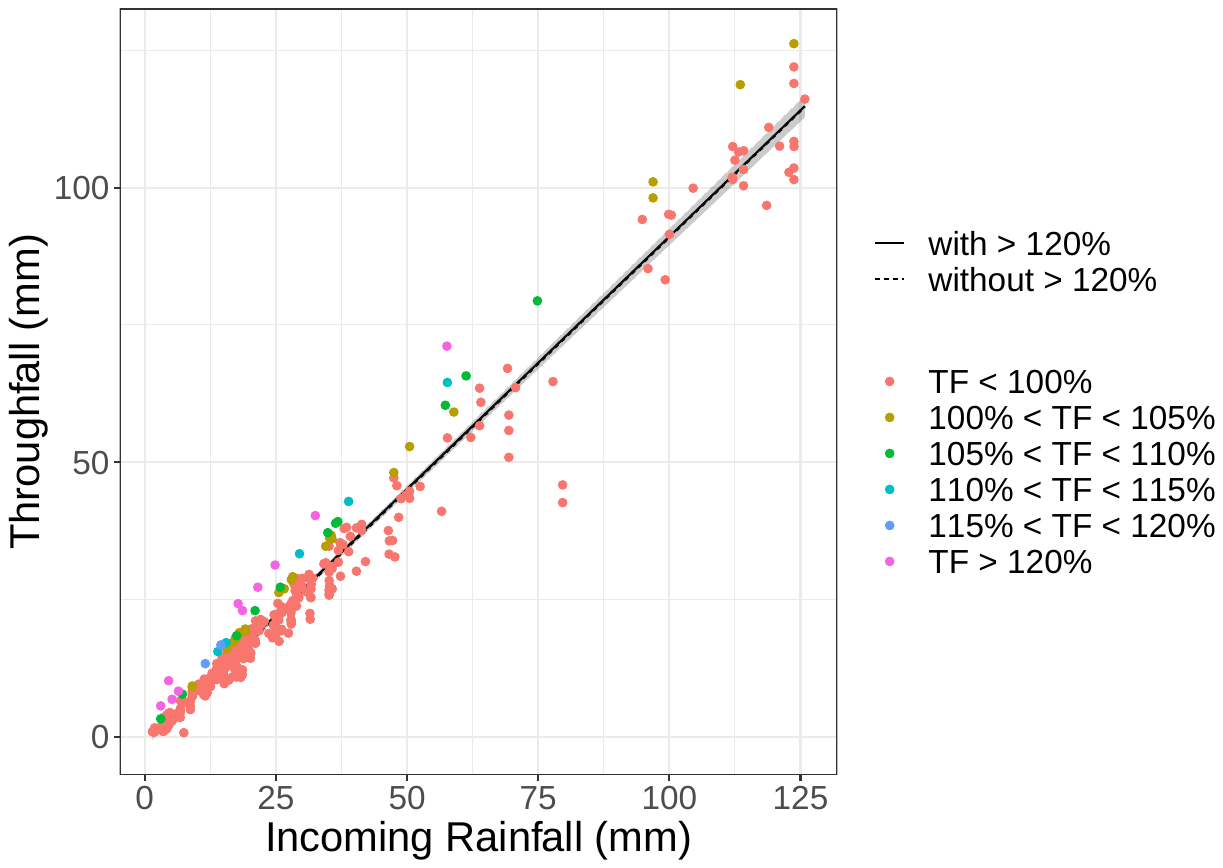

Figure S4: Relationship between average throughfall (mm) and incoming rainfall (mm), with the regression line based on all data (black line) and after excluding the measurements for which throughfall was >120% of incoming rainfall (purple symbols; dashed line)

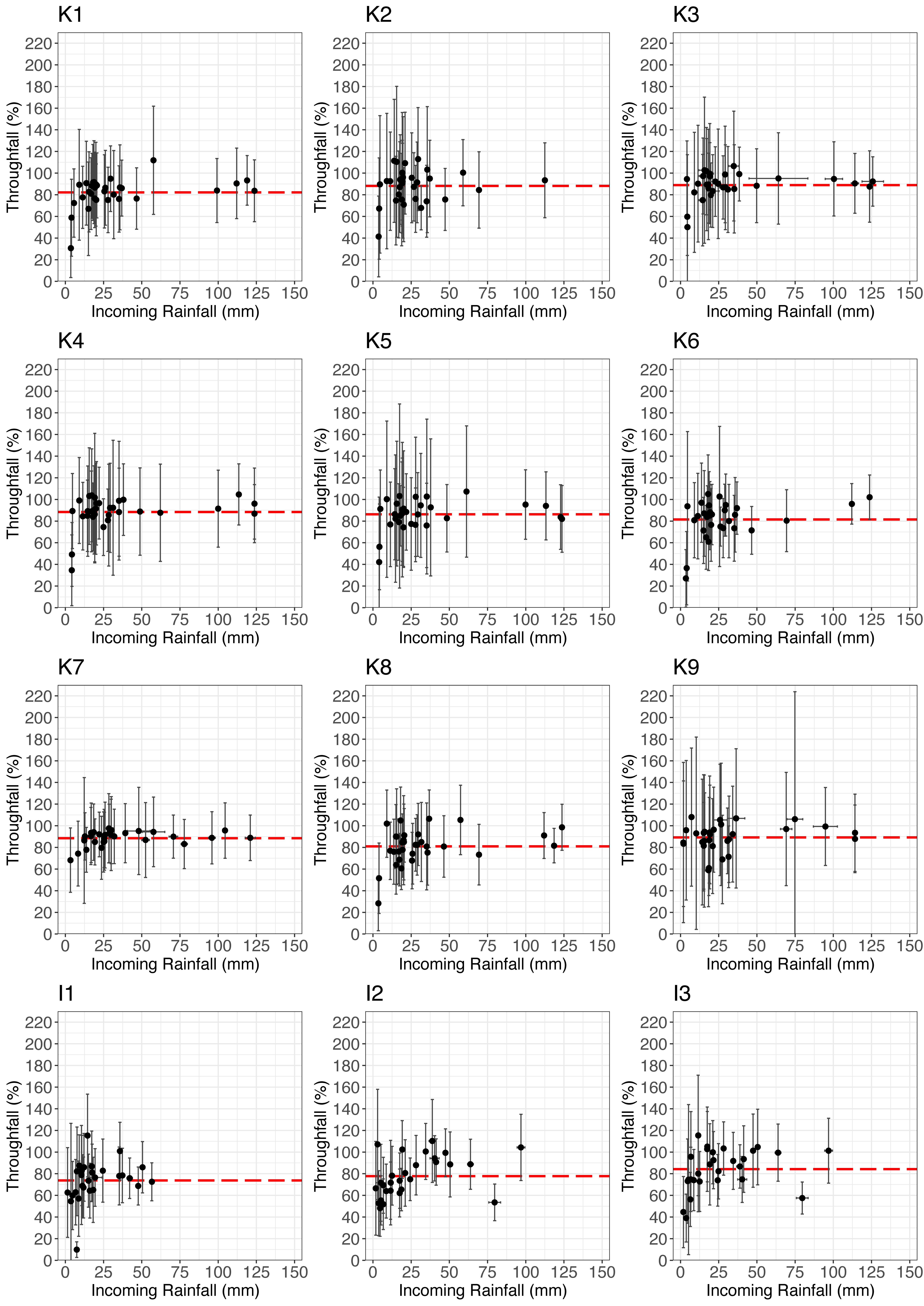

Figure S5: Relation between average throughfall (in % of reference rainfall) and incoming rainfall (mm) for each measurement day (n = 28 to 32) for the twelve forest plots (K1 – K9 (located in Kuamut area), and I1 – I3 (located in INFAPRO area). The symbol represents the average and the black, vertical line the standard deviation for the 50 measurements. The horizontal error bars represent the standard deviation of the reference rainfall measurements (only for plots with nearby reference rainfall stations: K3, K7, K9, I1, I2, I3). Please note, that in some case the standard deviation of the reference rainfall was very small and error bars are therefore not visible. The red dashed line shows the mean throughfall for each plot.

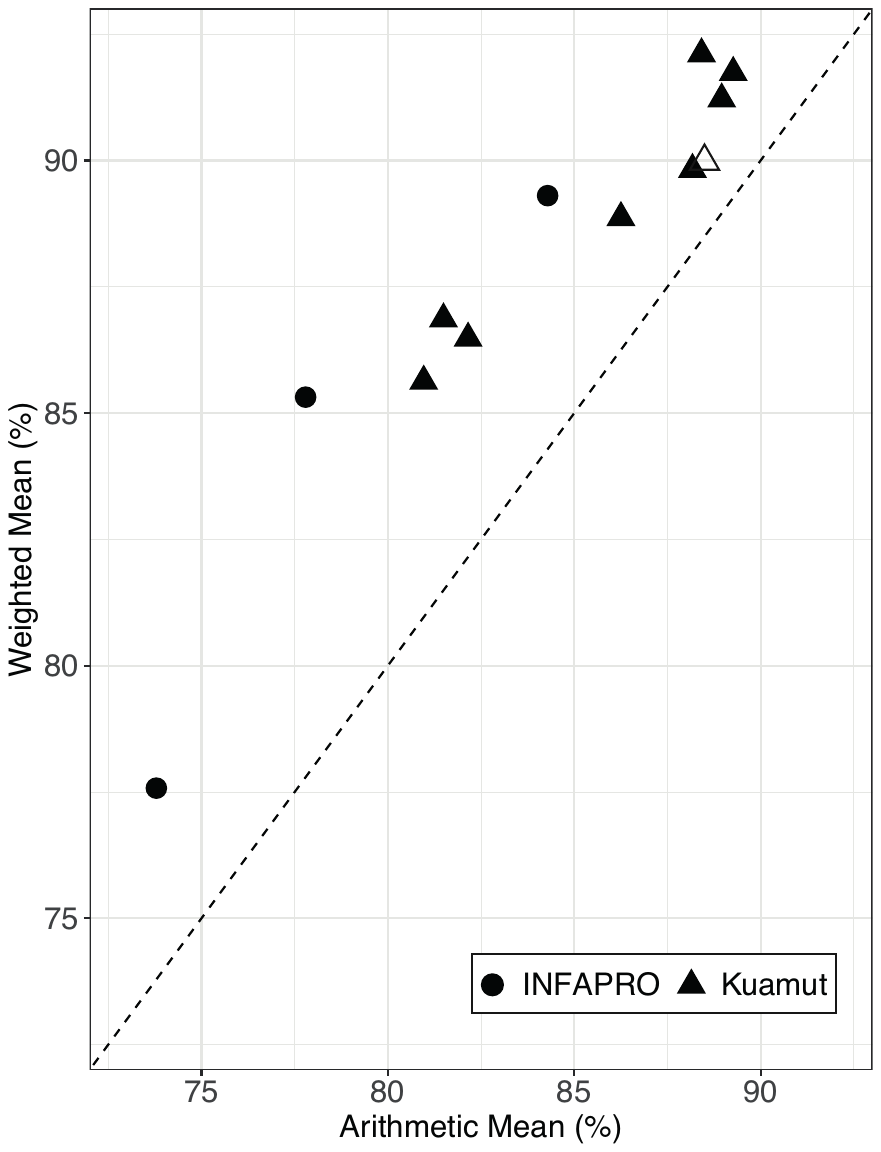

Figure S6: Relation between the arithmetic mean throughfall (in % of incoming rainfall) and the mean throughfall weighted by the incoming rainfall event size (also in % of incoming rainfall) for the 12 study plots. The plots in Kuamut are indicated by a triangle and the plots at INFAPRO by a circle. Forest plot K7 with the notably low Shannon Diversity Index is indicated by an open triangle (cf. Figure 2). The dashed line shows the 1:1 line.

**Supplementary material 4: Additional information on best fitting models**

Table S3: Linear model coefficients (n (i.e., number of observations), p-value, AIC, estimates of intercept, F-statistics, Adj. R-squared) of linear regression models for the relation between either ground-derived or LiDAR-derived diversity indicators and mean throughfall. The linear models with the Shannon Diversity Index are reported for two subsets of data (with and without plot K7; n=12 / n = 11). Please note, that the Aikake information criterion (AIC) is only comparable if n is congruent. Comparisons can be made with adjusted AIC values reported in Table S4.

| **ground-derived** | **Shannon Diversity Index** | n | 12 / 11 |
| --- | --- | --- | --- |
|  |  | p-value | 0.14 / 0.05 |
|  |  | AIC | 75.0 / 66.8 |
|  |  | Estimate (Intercept) | 93.37 / 121.89 |
|  |  | Estimate (Shannon Diversity Index) | -2.86 / -11.10 |
|  |  | F-statistics | 2.534 on 1 and 10 DF /  5.15 on 1 and 9 DF |
|  |  | Adj. R-squared | 0.12 / 0.29 |
|  | **Coefficient of variation (CV) of DBH** | n | 12 |
|  |  | p-value | 0.05 |
|  |  | AIC | 72.7 |
|  |  | Estimate (Intercept) | 98.3 |
|  |  | Estimate (CV of DBH) | -17.42 |
|  |  | F-statistics | 5.129 on 1 and 10 DF |
|  |  | Adj. R-squared | 0.27 |
| **LiDAR-derived** | **Coefficient of variation (CV) of Top of Canopy Height (TCH)** | n | 9 |
|  |  | p-value | 0.37 |
|  |  | AIC | 51.8 |
|  |  | Estimate (Intercept) | 74.84 |
|  |  | Estimate (CV of TCH) | 22.2 |
|  |  | F-statistics | 3.50 on 1 and 7 DF |
|  |  | Adj. R-squared | -0.01 |

Table S4: Model coefficients of simple and multiple linear regression models for different subsets of data. n gives the number of data points (=plots) used in the model, as well as the p-value, intercept, the estimate of main effects, F-statistics, Adj. R-squared, and the AIC values for these models. None of the interaction terms were significant. For n=11 data of forest plot K7 is excluded, for n = 9 data from forest plots I1, I2, and I3 are excluded, and for n = 8 data from forest plots K7, I1, I2, I3 was excluded. Grey-shaded boxes indicate the models that are discussed in section 4.4.

|  | **ground-derived** | | | | | | | | | | | | | **LiDAR-derived** | | | |
| --- | --- | --- | --- | --- | --- | --- | --- | --- | --- | --- | --- | --- | --- | --- | --- | --- | --- |
|  | **forest characteristic only** | | **With diversity indicator** | | | | | | | | | | | **forest characteristic only** | **With diversity indicator** | | |
|  |  | | Shannon Diversity Index (SDI) | | | | | | CV of DBH | | | | |  |  | | |
|  | tree density | tree height | tree density | SDI | | tree height | | SDI | tree density | CV of DBH | tree height | CV of DBH | | TCH | TCH | | CV of TCH |
| N | 12 | 12 | 12 | | | 12 | | | 12 | | 12 | | | NA | NA | | |
| p-value | 0.003 | 0.92 | 0.01 | 0.75 | | 0.96 | | 0.17 | 0.01 | 0.15 | 0.76 | 0.06 | | NA | NA | | NA |
| Intercept | 100.65 | 85.35 | 101.44 | | | 92.91 | | | 105.66 | | 95.43 | | | NA | NA | | |
| Estimate  (main effects) | -0.04 | -0.07 | -0.04 | -0.52 | | 0.03 | | -2.87 | -0.04 | -9.59 | 0.19 | -17.95 | | NA | NA | | NA |
| F-statistics | 15.05 on 1 and 10 DF | 0.01 on 1 and 10 DF | 6.91 on 2 and 9 DF | | | 1.14 on 2 and 9 DF | | | 9.86 on 2 and 9 DF | | 2.39 on 2 and 9 DF | | | NA | NA | | |
| Adj. R-squared | 0.56 | -0.10 | 0.52 | | | 0.03 | | | 0.62 | | 0.20 | | | NA | NA | | |
| AIC* | 66.7 | 77.7 | 68.5 | | | 77.0 | | | 65.7 | | 74.6 | | | NA | NA | | |
| N | 11 | 11 | 11 | | | 11 | | | 11 | | 11 | | | 11 | 11 | | |
| p-value | 0.007 | 0.88 | 0.03 | 0.17 | | 0.33 | | 0.04 | 0.01 | 0.12 | 0.76 | 0.11 | |  |  | | |
| Intercept | 101.04 | 85.61 | 119.43 | | | 120.19 | | | 108.61 | | 95.64 | | | NA | NA | | |
| Estimate (main effects) | -0.04 | -0.11 | -0.04 | -6.34 | | 0.65 | | -13.85 | -0.04 | -11.73 | 0.20 | -18.37 | | NA | NA | | NA |
| F-statistics | 11.95 on 1 and 9 DF | 0.02 on 1 and 9 DF | 8.027 on 2 and 8 DF | | | 3.12 on 2 and 8 DF | | | 8.87 on 2 and 8 DF | | 1.67 on 2 and 8 DF | | | NA | NA | | |
| Adj. R-squared | 0.52 | -0.11 | 0.58 | | | 0.30 | | | 0.61 | | 0.12 | | | NA | NA | | |
| AIC**^+^** | 62.5 | 71.8 | 61.7 | | | 67.5 | | | 61.0 | | 70.0 | | | NA | NA | | |
| N | 9 | 9 | 9 | | | 9 | | | 9 | | 9 | | | 9 | 9 | | |
| p-value | 0.02 | 0.11 | 0.04 | 0.95 | | 0.14 | | 0.50 | 0.03 | 0.19 | 0.07 | 0.10 | | 0.18 | 0.28 | 0.56 | |
| Intercept | 96.9 | 71.79 | 97.05 | | | 75.34 | | | 100.99 | | 79.63 | | | 91.62 | 83.58 | | |
| Estimate (main effects) | -0.03 | 0.86 | -0.03 | -0.08 | | 0.82 | | -0.94 | -0.03 | -6.84 | 0.86 | -10.00 | | -0.23 | -0.20 | | 14.71 |
| F-statistics | 9.30 on 1 and 7 DF | 3.40 on 1 and 7 DF | 2.22 on 3 and 5 DF | | | 1.84 on 2 and 6 DF | | | 6.54 on 2 and 6 DF | | 4.29 on 2 and 6 DF | | | 2.2 on 1 and 7 DF | 1.198 on 2 and 6 DF | | |
| Adj. R-squared | 0.51 | 0.23 | 0.31 | | | 0.17 | | | 0.58 | | 0.45 | | | 0.13 | 0.04 | | |
| AIC**^±^** | 45.3 | 49.4 | 49.3 | | | 50.6 | | | 44.50 | | 46.9 | | | 50.5 | 51.9 | | |
| N | 8 | 8 | 8 | | | 8 | | | 8 | | 8 | | | 8 | 8 | | |
| p-value | 0.04 | 0.15 | 0.06 | | 0.73 | 0.15 | 0.47 | | 0.04 | 0.17 | 0.10 | | 0.15 | 0.29 | 0.31 | 0.55 | |
| Intercept | 96.99 | 72.34 | 102.61 | | | 85.24 | | | 103.2 | | 80.01 | | | 91.26 | 81.22 | | |
| Estimate (main effects) | -0.03 | 0.81 | -0.03 | | -1.72 | 0.90 | -4.31 | | -0.03 | -8.4 | 0.88 | | -10.87 | -0.22 | -0.23 | 20.922 | |
| F-statistics | 6.98 on 1 and 6 DF | 2.65 on 1 and 6 DF | 3.06 on 2 and 5 DF | | | 1.53 on 2 and 5 DF | | | 5.615 on 2 and 5 DF | | 3.237 on 2 and 5 DF | | | 1.347 on 1 and 6 DF | 0.8065 on 2 and 5 DF | | |
| Adj. R-squared | 0.46 | 0.19 | 0.37 | | | 0.13 | | | 0.57 | | 0.39 | | | 0.05 | -0.06 | | |
| AIC**^×^** | 41.9 | 45.1 | 43.7 | | | 46.2 | | | 40.6 | | 43.4 | | | 46.4 | 47.8 | | |
